# Unisexual flower initiation in the monoecious *Quercus suber* L., a molecular approach

**DOI:** 10.1101/2020.02.13.945295

**Authors:** Rómulo Sobral, Helena Gomes Silva, Sara Laranjeira, Joana Magalhães, Luís Andrade, Ana Teresa Alhinho, Maria Manuela Ribeiro Costa

## Abstract

Several plant species display a temporal separation of the male and female flower organ development to enhance outbreeding, however, little is known regarding the genetic mechanisms controlling this temporal separation. *Quercus suber* is a monoecious oak tree with accentuated protandry: in late winter, unisexual male flowers emerge adjacent to the swollen buds, whereas unisexual female flowers emerge in the axils of newly formed leaves formed during spring (4 to 8 weeks after male flowering). Here, phylogenetic profiling lead to the identification of cork oak homologs of key floral regulatory genes. The role of these cork oak homologs during flower development was identified with functional studies in *Arabidopsis thaliana*. The expression profile of flower regulators (inducers and repressors) throughout the year, in leaves and buds, suggests that the development of male and female flowers may be preceded by separated induction events. Female flowers are most likely induced during the vegetative flush occurring in spring, whereas male flowers may be induced in early summer, staying enclosed within the pre-dormant buds, but complete their development before the vegetative flush of the following year, displaying a long period of anthesis that spans the dormant period. Results portray a genetic mechanism that may explain similar reproductive habits in other tree species.

## Introduction

Temporal separation of male and female flowering events (dichogamy) is a usual observed feature in perennial species and a mechanism that has emerged independently during evolution to avoid the adverse effects of inbreeding (Polito and Li 1985, Pendleton et al. 2000, Bai et al. 2007, Renner et al. 2007, Gleiser et al. 2008, Watanabe et al. 2016). The timing of male and female organ development in dichogamic species can vary quite substantially, from 6 hours, in the case of the hermaphrodite *Annona squamosa* (Wester 1910), to weeks in the case of the monoecious *Carya illinoiensis* (Shuhart 1927, Woodroof 1927). Therefore, it is possible that in species such as the latter, development of the male and female unisexual flowers may depend on distinct flowering induction events. Despite the economic importance of some dichogamic species, little is known about the genetic mechanisms controlling the initiation of the independent flower reproductive organs.

The regulatory mechanisms controlling flower induction have been extensively studied in annual model species (reviewed in Wang, 2014; Capovilla et al., 2015; Johansson and Staiger, 2015). Contrary to annual species, perennials are required to coordinate multiple flowering events during their lifespan that imply additional regulation tightly controlled by seasonal cues (reviewed in Samach and Smith, 2013). As an example, in temperate perennial species, the annual growth cycle is characterized by vegetative/reproductive growth during spring followed by a dormant phase during the cold season that protects meristems from unfavourable environmental conditions, thus sustaining the perennial habit (Rohde et al. 2000). However, despite the intrinsic differences between annual and perennial life cycles, recent new tools in tree genomics suggest that the bulk of the flowering induction machinery identified in annuals is most likely conserved in perennials, and that species-specific reproductive features may be correlated with specific perturbations in conserved regulatory networks (Böhlenius et al. 2006, Hsu et al. 2006, 2011, Ibáñez et al. 2010, Varkonyi-Gasic et al. 2011, Karlberg et al. 2011, Neale and Kremer 2011, Karlgren et al. 2013, Azeez et al. 2014, Tylewicz et al. 2015, Zhang et al. 2016, Voogd et al. 2017, Miyazaki and Satake 2017, Endo et al. 2018, Satake, Kawatsu, Chiba, et al. 2019, Satake, Kawatsu, Teshima, et al. 2019).

Regulation of flower induction, both in annual and perennial species, is dependent on three distinct pathways: photoperiod/light perception, temperature and age (reviewed in Wang, 2014; Capovilla *et al*., 2015; Johansson and Staiger, 2015). The age pathway ensures that juvenile plants, that have not enough biomass to support reproduction, do not flower even if subjected to flowering inductive cues. The age pathway is controlled by microRNA156 (miR156) and its targets, the *SQUAMOSA PROMOTER BINDING-LIKE* (*SPL*s) genes, in both annual and perennial plants (Wang et al. 2011, Wang 2014). The expression of *miR156* is high in young plants and steadily decreases as plants age (Wu and Poethig 2006, Wu et al. 2009). As a consequence, in the juvenile phase, the levels of miR156-targeted *SPL*s transcripts are low. As the plants age, the amount of miR156 decreases, resulting in an increasing stability of miR156-targeted *SPL*s. In the model species *Arabidopsis thaliana*, *SPL*s promote flowering by increasing the expression of *miR172* and flower regulators, such as, *FLOWERING LOCUS T* (*FT*), *FRUITFUL* (*FUL*), and *APETALA1* (*AP1*) (Wang, Czech, et al. 2009, Wu et al. 2009, Yamaguchi et al. 2009, Wang et al. 2011).

Light and its duration (photoperiod) over the course of a day are important cues that control flower induction. By slightly different mechanisms, some species respond to long days (LD) and others to short days (SD). Whichever mechanism is used to respond to different photoperiods, the main output of the interaction between the circadian clock and the light cues is the regulation of a *CONSTANS*-like factor that tightly control the expression of a *FT-*like gene, whose gene product is commonly known as the florigen (reviewed in Golembeski *et al*., 2014; Song *et al*., 2015). The ectopic expression or down-regulation of *FT*-like genes in a broad range of plant species have demonstrated that *FT*-like genes act as universal promoters of flowering (Endo et al. 2005, Böhlenius et al. 2006, Hsu et al. 2006, 2011, Tränkner et al. 2010, Fu et al. 2014, Ye et al. 2014, Freiman et al. 2015, Klocko et al. 2016).

The *FT* locus is a checkpoint for other flowering pathways (*e.g.* temperature) and, as such, is a direct regulatory target of several floral repressors including *FLOWERING LOCUS C (FLC)* and *SHORT VEGETATIVE PHASE* (*SVP*), that are the canonical output of the temperature/vernalization pathway by down-regulating *FT* and *SUPRESSOR OF OVEREXPRESSION OF CONSTANS 1* (*SOC1*) (Michaels and Amasino 1999, Hartmann et al. 2000, Sheldon et al. 2000, Li et al. 2008, Tao et al. 2012, Hwan Lee et al. 2013, Posé et al. 2013). In trees, *DORMANCY-ASSOCIATED MADS-BOX*-like genes, which are *SVP* homologs, have been proposed to control bud dormancy establishment and are distinctly affected by photoperiod and temperature signals (Li et al. 2009, Horvath et al. 2010, Jiménez et al. 2010, Yamane et al. 2011, Saito et al. 2013, Howe et al. 2015).

In Arabidopsis, FT interacts with FD in the shoot apical meristem, (Abe et al. 2005, Wigge et al. 2005, Taoka et al. 2011) triggering a transcriptional machinery that activate several downstream transcription factor genes such as *AP1*, *FUL*, *SOC1* and *LEAFY* (*LFY*) committing the apical meristem into developing flowers (Abe et al. 2005, Wigge et al. 2005). *LFY* belongs to a highly conserved gene family that share an important role in the establishment of floral meristems and floral organ identities in several species across the angiosperms (Blázquez et al. 1997, Rottmann et al. 2000, Peña et al. 2001). *LFY* is the primary activator of the floral organ identity genes and directly activates *AP1* (flower meristem identity gene/ A-class gene, necessary to produce sepals and petals), *APETALA3* and *PISTILLATA* (B-class genes, necessary to the development of petals and stamens), *AGAMOUS* (*AG*) (C-class gene, necessary to stamens and carpel development) and *SEPALLATA3* (SEP3) (E-class gene, necessary to the development of petals, stamens and carpels) (Bowman et al. 1989, 1993, Coen and Meyerowitz 1991, Weigel and Nilsson 1995, Flanagan et al. 1996, Parcy et al. 1998, Liljegren et al. 2000, Pelaz et al. 2000, Moyroud et al. 2011, Winter et al. 2011).

A considerable lag of time between the onset of the male and female functions is frequent in woody tree species across distinct angiosperm taxa, such as *C. illinoiensis* (pecan), *Castanea sativa* (chestnut), *Juglans regia* (walnut), *Corylus avellana* (hazelnut), Quercus *spp*. or Betula *spp*. (reviewed in Sedgley and Griffin, 1989). *Quercus suber* is a perennial species abundant in the Mediterranean climate zone, particularly in Southern Europe (Portugal, Spain, France and Italy) and Northern Africa (Morocco, Northern Algeria and Tunisia). Cork oak trees have an extended juvenile phase before they start producing flowers (approximately 15 years). The vegetative growth has a clear temperate seasonality, with a phase of active growth and a phase of bud dormancy. Active growth starts usually in March with the emergence of new leaves, and continues until late summer/early autumn, when growth ceases and the tree enters a period of dormancy until the following spring (Natividade 1950, Elena-Rossello et al. 1993, Oliveira et al. 1994, Varela and Valdiviesso 1996, Gómez-Casero et al. 2007). In common with other Quercus species, *Q. suber* is a monoecious tree with a pronounced protandrous system (male flowering precedes female flowering) (Natividade 1950, Ledig et al. 1971, Tisserat et al. 1979, Kaul 1985). In late winter, male flowers (organized in catkins) emerge adjacent to the buds formed during the previous growth season. Female flowers appear in the axils of newly formed leaves during spring after the vegetative flush, four to eight weeks after male flowering. Occasionally, a male flower bloom (but never female flowering) occurs in autumn during the establishment of dormancy (Varela and Valdiviesso 1996, Boavida et al. 1999).

Here, a functional analysis of the *Q. suber* homologs of key floral regulatory genes (inducers and repressors) was conducted in order to understand the mechanisms that control the protandric habit of *Q. suber*. To identify events leading to flower induction, expression of the cork oak flower induction homologs was analysed in leaves and buds throughout the year, including the growth and dormancy seasons. Results suggest that an induction event occurring in summer is likely to be associated with the formation of male unisexual flowers that stay dormant during winter and only resume their development during bud swelling in the following spring, coinciding with a second event of flower induction that leads to the formation of female flowers. Our results shed a new light on the molecular events that control the protandrous habit in cork oak and may also explain similar reproductive habits in other tree species, including other oaks and other species with high agronomic interest.

## Material and Methods

### Plant material

*Arabidopsis thaliana* ecotype Columbia seeds (Col-0) were acquired from the Nottingham Arabidopsis Stock Centre. Seeds were sown in Murashige and Skoog agar medium and were incubated under long-day conditions (16 hours light/8 hours dark) at 20 °C in controlled environmental growth rooms, with light intensity of 70 μ E m^-2^ s^-1^. Approximately 10 days after sowing, plantlets were transferred to pots containing a 4:1 (v:v) mixture of turf rich soil and vermiculite. Overexpression constructs were introduced into *A. thaliana* ecotype *Columbia* using the GV3101 Agrobacterium strain and the floral dip method (Clough and Bent 1998). Transgenic plants were selected in medium supplemented with hygromiycin (50 µg/mL). After checking for single insertion lines, four independent lines were selected for each construct. Functional studies were conducted using T3 homozygous lines (with a single insertion locus, segregation 3:1 in the T2 generation). Three *Q. suber* adult trees, located at the Minho University campus, were chosen based on similar phenological characteristics, such as timing of axillary bud growth cessation, axillary bud swelling, leaf initiation, leaf development and male and female flower initiation across four recorded growth/dormancy seasons. The juvenile tree was selected due to its smaller size and inability to flower. Phenological observations on the juvenile tree showed that the periods/stages of axillary bud growth cessation, axillary bud swelling, leaf initiation and leaf development were coincident with the ones recorded from the adult trees. Plant organ sampling (axillary buds, leaves, male and female flowers) was conducted in three consecutive years twice a month. Expression data from the third year sampling was excluded from any analysis due to consecutive days of abnormally high temperatures during the growth cessation period.

Samples were collected from southern facing branches in the morning, approximately 2-4 hours after sunrise. Light intensity during collection was not recorded. In each sampling, 3-5 buds were randomly collected from different positions of the branch. Similarly, each leaf sampled was chosen randomly within the same branch. Male and female inflorescences were collected based on the phenological stages defined by Varela and Valdiviesso (1996). Flowering time in Arabidopsis plants was measured by counting the total number of leaves in each plant (n=20) (Suárez-López et al. 2001). Flowering time measurements were repeated three times independently. A student t-test was performed to statistically differentiate the wild type and the transgenic lines.

### Phylogenetic methods

Cork oak protein sequences were obtained by performing a BLAST in the cork oak database (www.corkoakdb.org) and the *Q. suber* genome database (www.ncbi.nlm.nih.gov/) using the *A. thaliana* protein sequences as query (Table S1). Homologous sequences from other selected species were obtained by performing a PSI-BLAST at the NCBI (www.ncbi.nlm.nih.gov/), PLAZA databases (https://bioinformatics.psb.ugent.be/plaza/) and *Q. robur* transcriptome and genome (https://urgi.versailles.inra.fr/blast). Protein sequences were aligned with CLUSTALW (Löytynoja and Goldman 2005). Distances were estimated using the Jones-Taylor-Thornton (JTT) model of evolution for a maximum-likelihood tree with the MEGA6 software package (Felsenstein 1986). The trees were rooted using the outgroup as the root. To provide statistical support for each node on the tree, a consensus tree was generated from 1000 bootstrap data sets.

### RNA extraction and cDNA preparation

*Arabidopsis thaliana* total RNA was extracted using TRIzol^TM^ reagent (Termo Fisher Scientific) according to the manufacturer’s instructions. *Quercus suber* total RNA was obtained using the CTAB/LiCl extraction method (Chang et al. 1993) with some modifications (Azevedo et al. 2003)*. A. thaliana* and *Q. suber* cDNA were synthesized according to the Invitrogen cDNA synthesis kit SuperScript® III RT manufacturer’s instructions. *Q. suber miRNA* expression was analysed using a cDNA that was synthesized with the Invitrogen NCode™ miRNA qRT-PCR kit.

### qRT-PCR analysis

cDNA was amplified using SsoFast™ EvaGreen® Supermix (Bio-Rad), 250 nM of each gene-specific primer (listed in Table S2) and 1 μL of cDNA (1:100 dilution). qRT-PCR reactions were performed in triplicates on the CFX96 Touch™ Real-Time PCR Detection System (Bio-Rad). After an initial period of 3 min at 95 °C, each of the 40 PCR cycles consisted of a denaturation step of 10 s at 95 °C and an annealing/extension step of 10 s at the gene specific primer temperature. With each PCR reaction, a melting curve was obtained to check for amplification specificity and reaction contamination, by heating the amplification products from 60 °C to 95 °C in 5 s intervals. Primer efficiency was analysed with CFX Manager™ Software v3.1 (Bio-Rad), using the Livak calculation method for normalized expression (Livak and Schmittgen 2001). Gene expression analysis was established based on three technical of three adult and one juvenile trees, and normalized with the reference genes *QsPP2AA3* or *QsACTIN* (Marum et al. 2012).

### Plasmids construction

Overexpression constructs were obtained using gateway technology (Thermo Fisher Scientific) according to the manufacturer’s instructions. The destination vector was pMDC32. All the primers used are described in Table S2.

## Results

### Reproductive development of *Q. suber* follows a temperate habit

*Quercus suber* is a monoecious tree species that exhibits a typical temperate perennial habit. During autumn and possibly in response to shorter days and/or lower temperatures, the shoot meristems cease activity and enter dormancy (while enclosed in a bud structure) (Fig. 1a and 1b) similarly to what has been described for other Quercus species (Derory et al. 2006). The axillary and terminal buds stay dormant throughout winter probably due to the persistency of low temperatures (Fig. 1c, black arrow). In late February/early March, the increase in day length and/or rising temperatures lead to the resume of meristematic activity and, thus, to bud swelling, signalling the end of the dormancy period (Fig. 1d and Fig. 1e, asterisks). In adult trees, male flowers with well-defined anthers emerge in the axils of the swelling axillary buds (Fig. 1d and 1e, white arrows).

**Figure 1.**
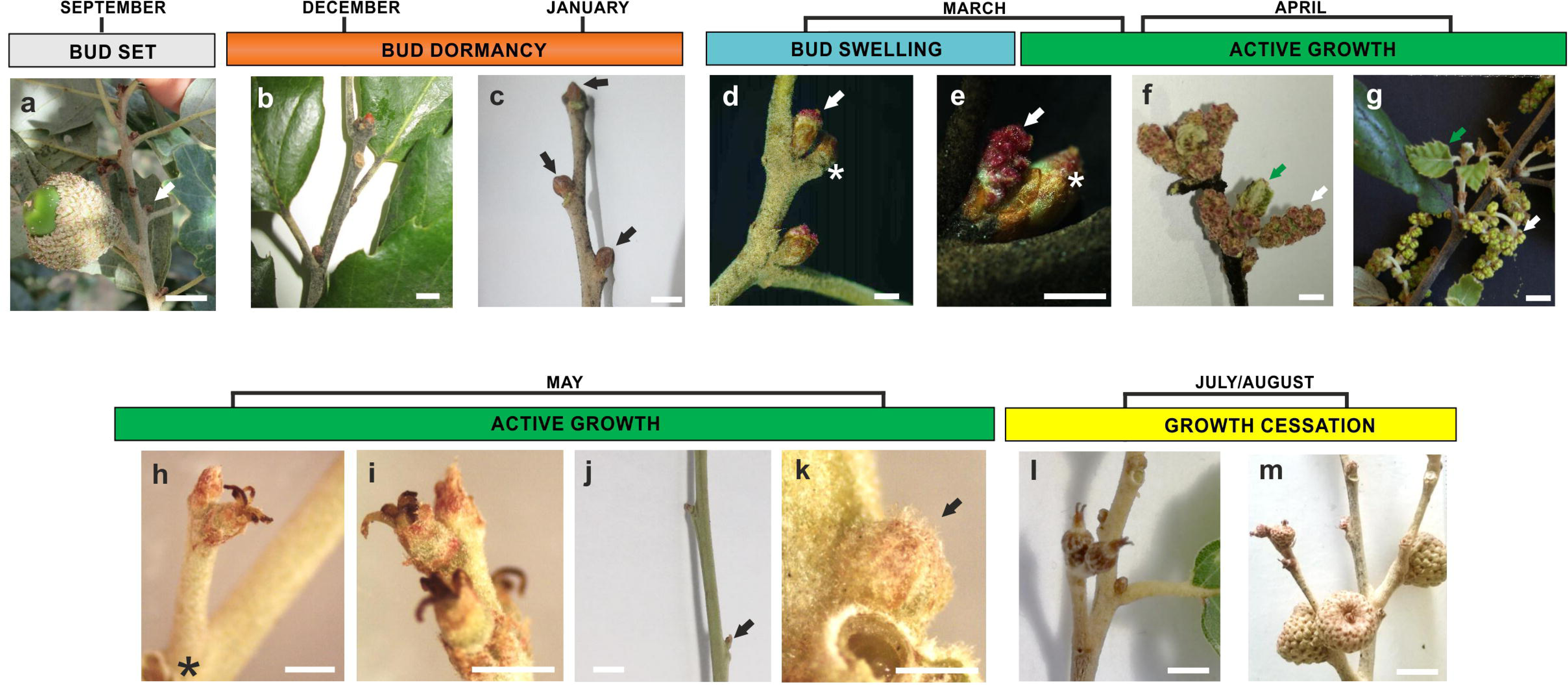
*Quercus suber* vegetative and reproductive features throughout the year. **a** - *Q. suber* branch displaying an axillary bud during growth cessation in September (arrow indicates an axillary bud). **b** - *Q. suber* branch containing arrested axillary buds in December. **c** - *Q. suber* branch containing arrested axillary buds (black arrows – axillary buds) collected in January (axillary buds are located in the axil of leaves that were removed to improve the quality of the picture). **d** - Male flowers emerge adjacent to the swelling axillary bud (white arrow – male inflorescence, white asterisk – swelling bud) in March. **e** - Male flowers develop in the axil of the swelling axillary bud (white arrow – male inflorescence, white asterisk – swelling bud) in March. **f** - In April, male flower completes development (white arrow) and vegetative flush occurs (green arrow) after axillary bud burst. **g** - In late April, male flowers become mature while new leaves develop**. h** - In May, female inflorescences develop in the axil of new leaves (leaves were removed to improve the quality of the picture). Black asterisk marks the leaf position. **i** - In May, female spike contains female flowers at distinct developmental stages. **j** - Small axillary buds (black arrow) are formed in the axils of new leaves that do not show female flowers (leaves subtending the axillary buds were removed to improve the quality of the picture). **k** - Magnification of the image containing the axillary bud depicted in J. Black arrow represents a small axillary bud. **l** and **m** - Female flowers develop into acorns in July/August. Scale: 0.5 mm.

In April, male flowers complete their development (Fig. 1f and 1g, white arrows), the swollen axillary buds burst open and new shoots emerge containing young leaves (Fig. 1f and 1g, green arrows). Occasionally, an abnormal burst of unisexual male flowers (but never of female flowers) occurs in September/October, at a time when dormancy is normally being established. This occurs in some trees when subjected to abnormal warm autumn temperatures. Female flowers are observed in the new shoots in April and May, 4 to 8 weeks after the emergence of the male flowers. Female flowers are organized in spikes of 1-8 carpellate flowers and develop in the axils of the newly formed leaves (Fig. 1h (asterisk) and 1i). Female flowers within the same spike develop in an acropetal manner, which is suggestive of an active inflorescence meristem able to produce flowers for a short period of time. Female flowers stop developing during spring and later, in summer, the pollinated female flowers resume growth and develop into acorns (Fig. 1l and 1m).

In the axils of new leaves that do not have a female flower spike, there is the emergence of a small axillary bud structure (Fig. 1j and 1k, black arrows). Some of these new axillary buds will enlarge and enter dormancy in the autumn.

The morphological changes that occur during *Q. suber* reproductive development have been well described,(Varela and Valdiviesso 1996) but the mechanisms that control flower induction are still unknown in cork oak and in other Quercus species.

### Homologs of flowering time genes can be found in *Q. suber*

The control of flowering time is a complex process that relies on the balanced activity of regulators that promote or delay flowering until optimal conditions are met. Thus, to identify potential factors controlling the flower induction mechanism of male and female flowers of *Q. suber*, the cork oak transcriptome and genome databases were screened for homologs of *A. thaliana* gene hubs of distinct flower induction pathways (*FT*, *SOC1* and *SPLs*) (Kobayashi et al. 1999, Samach et al. 2000, Wu and Poethig 2006, Schwarz et al. 2008, Wang, Czech, et al. 2009). Flowering repressors that integrate the vernalization/temperature signalling pathways (*SVP* and *FLC*) (Michaels and Amasino 1999, Hartmann et al. 2000, Posé et al. 2013) were also used as queries in a BLAST search. All the genes identified in the cork oak genome BLAST search were also identified in the cork oak transcriptome BLAST search.

After retrieving the *Q. suber* homologs for the flowering regulator genes, a phylogenetic analysis was performed using available homologs from other perennial species (*Q. robur*, *Malus domestica*, *P. trichocarpa*, *Vitis vinifera, Prunus persica, Prunus mume*, *Actinidea chinensis, Fagus crenata, Citrus unshui, Eriobotrya deflexa, Euphorbia esula* and *Eucalyptus grandis*). Homologs with a known role in flowering induction were also included (*A. thaliana*, *Cardamine flexuosa*, *Antirrhinum majus*, *Solanum lycopersicum, Zea Mays, Oryza sativa, Arabis alpina*, *Petunia x hybrida, Glycine max, Medicago truncatula, Hordeum vulgare, Chrysanthemum seticuspe* and *Beta vulgaris*).

A phylogenetic tree was constructed using five *Q. suber* FT-like proteins and FT homologs from different species. The phylogenetic analysis also included protein sequences of gene homologs belonging to different members of the *FT* family, such as *TERMINAL FLOWER 1* (*TFL1*), *TWIN SISTER OF FT* (*TSF*), *MOTHER OF FT* (*MFT*) and *BROTHER OF FT* (*BFT*). The phylogenetic tree is divided into distinct clades, each one associated to a different sub-family (Fig. 2a). Two homologs belong to the TFL1 clade (QsTFL1 and QsCEN), another to the MFT clade (QsMFT), a fourth to the FT clade (QsFT) and the last homolog to the BFT clade (QsBFT) (Fig. 2a). The QsFT identified in the transcriptome of *Q. suber* is likely to be the closest homolog to AtFT.

**Figure 2.**
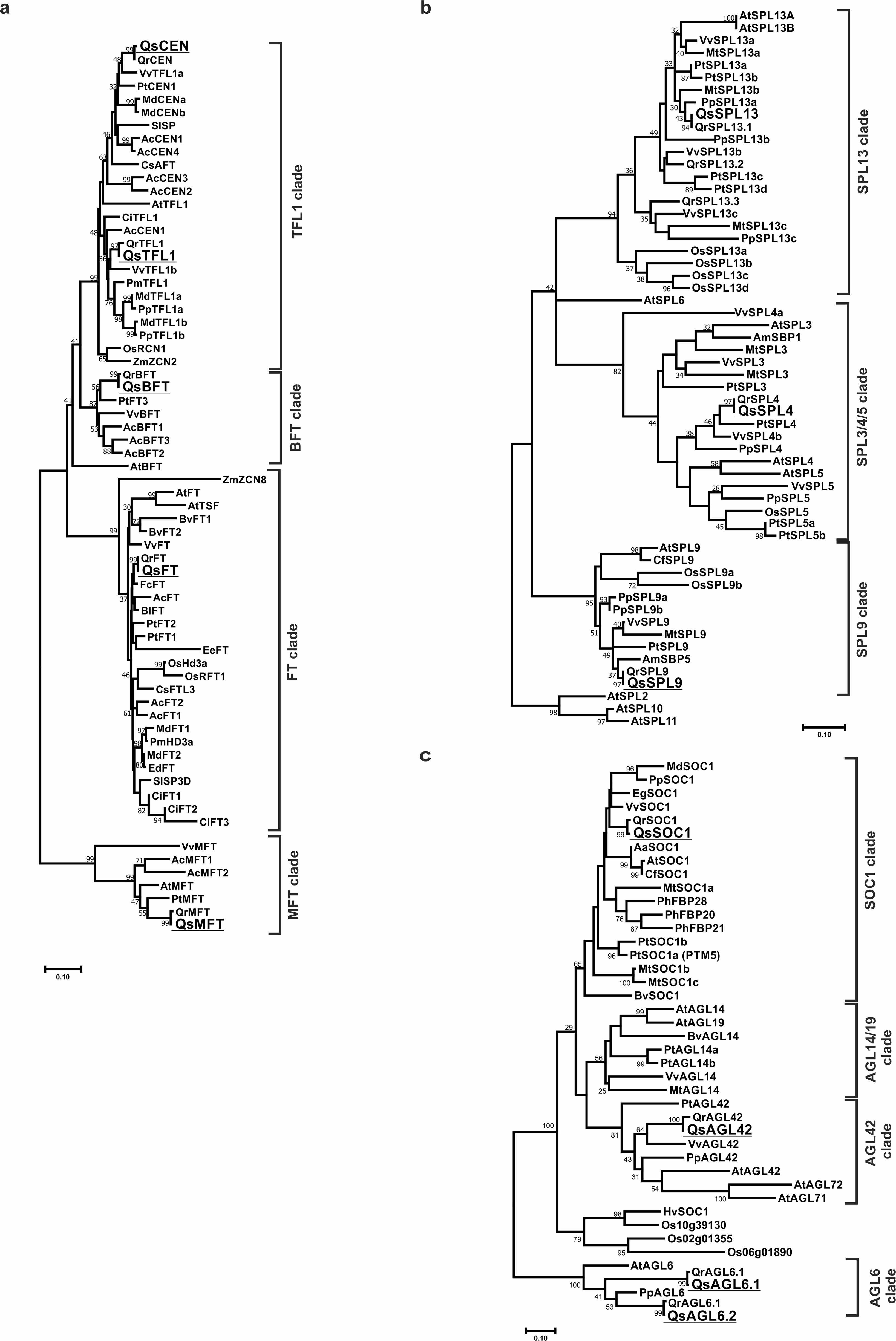
Homologs of positive regulators of floral induction are found in *Quercus suber*. **a** - FT family. **b -** SPL family. **c** - SOC1 family. The *Q. suber* proteins are underlined. The species used in the phylogenetic analysis were: *Quercus suber* (Qs), *Arabidopsis thaliana* (At), *Populus trichocarpa* (Pt), *Quercus robur* (Qr), *Vitis vinifera* (Vv), *Malus x domestica* (Md), *Solanum lycopersicum* (Sl), *Actinidea sitchensis* (Ac), *Oryza sativa* (Os), *Zea mays* (Zm), *Prunus persica* (Pp), *Beta vulgaris* (Bv), *Fagus crenata* (Fc), *Betula latifolia* (Bl), *Citrus sinensis* (Ci), *Chrysanthemum seticuspe* (Cs), *Euphorbia esula* (Ee), *Hordeum vulgare* (Hv), *Eriobotrya deflexa* (Ed), Prunus mume (Pm), *Medicago truncatula* (Mt), *Antirrhinum majus* (Am), *Cardamine flexuosa* (Cf), *Eucalyptus grandis* (Eg) and *Petunia hybrida* (Ph). Phylogenies were inferred using the Maximum-likelihood method. The percentage of replicate trees in which the associated taxa clustered together in the bootstrap test (1000 replicates) is shown next to the branches. The evolutionary distances were computed using the JTT matrix-based method and are in the units of the number of amino acid substitutions *per site*. Protein accession numbers used in the phylogenetic analysis are presented in table S1.

The SPL family is also characterized by extensive duplication as depicted in Figure 2b. The SPL family in Arabidopsis has fifteen members. Three SPL proteins were retrieved from the cork oak database and named after the respective homologs based on the phylogenetic analysis (QsSPL4, QsSPL9 and QsSPL13) (Fig. 2b).

According to Becker and Theißen (2003), the MADS*-*box SOC1 belongs to the TM3 sub-family together with five other members, AGAMOUS-LIKE 14 (AGL14), AGAMOUS-LIKE 19 (AGL19), AGAMOUS-LIKE 42 (AGL42), AGAMOUS-LIKE 71 (AGL71) and AGAMOUS-LIKE 72 (AGL72). By performing a BLAST search with the protein sequence of each member of the TM3 sub-family against the cork oak databases it was possible to identify two proteins. A phylogenetic analysis was then conducted using homologs for the different members of the TM3 clade of different species. AGAMOUS-LIKE6 clade was used as an out-group (Fig. 2c). The phylogenetic analysis suggests that one of the *Q. suber* proteins is homolog to SOC1, whereas the other is closer to AGL42 (Fig. 2c). No homologs were found for the other TM3 sub-family members in both *Q. suber* transcriptome or genome databases.

FLC together with FLOWERING LOCUS M (FLM), AT5G65070, AT5G65060, AT5G65080 and AGAMOUS-LIKE 31 (AGL31) belong to the FLC sub-family within the MADS family (Becker and Theißen 2003). Only one protein was retrieved when blasting the cork oak databases with the AtFLC protein sequence (QsFLC). The subsequent phylogenetic analysis indicated that QsFLC closely grouped with FLC-like proteins of other perennial species but was considerably distant from FLC-like proteins with a known role in flowering repression, such as FLC (AtFLC), *C. flexuosa* FLC (CfFLC) and *A. alpina* PEP1 (AaPEP1) (Fig. 3a) (Sheldon et al. 2000, Wang, Farrona, et al. 2009, Zhou et al. 2013).

**Figure 3.**
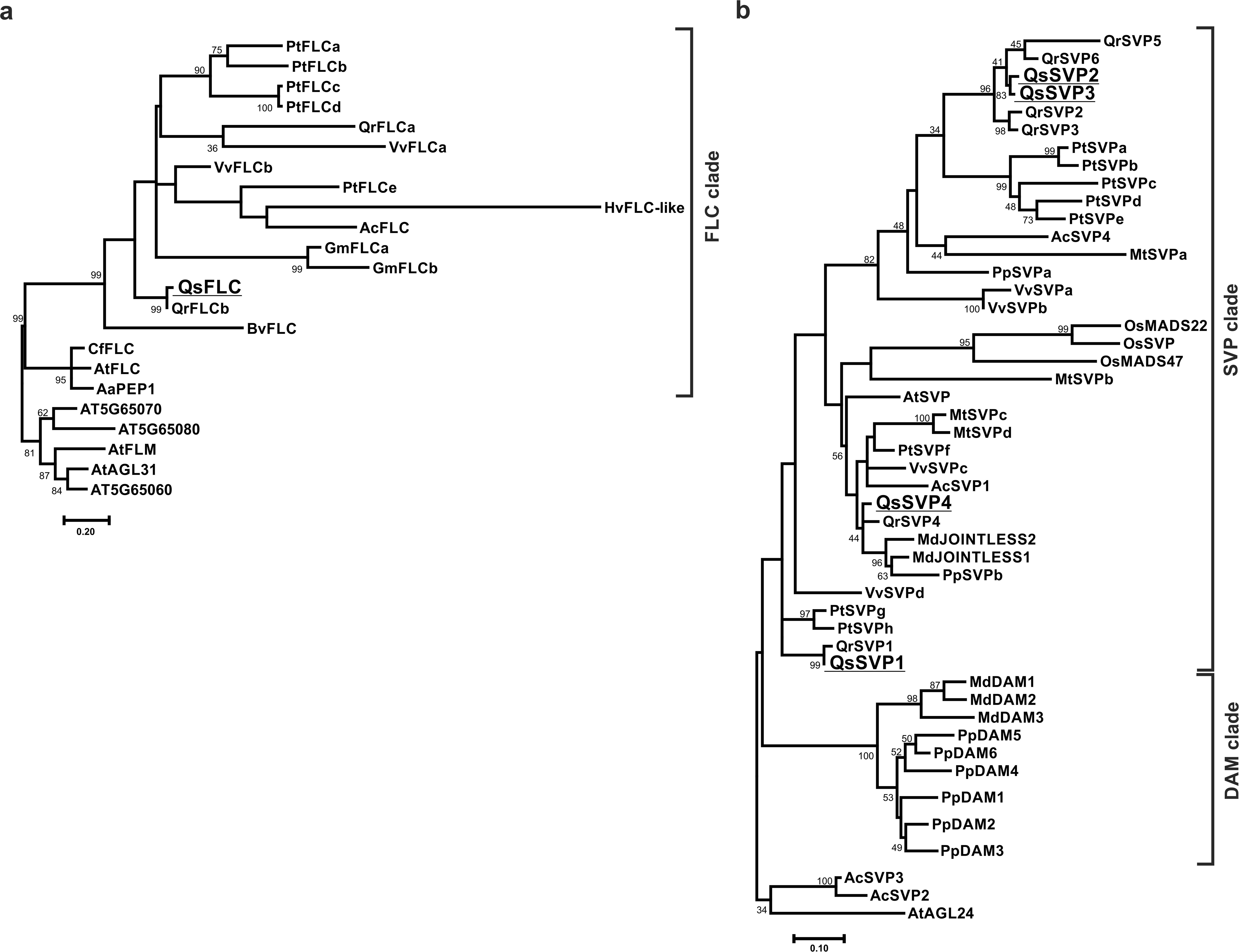
Homologs of negative regulators of floral induction are found in *Quercus suber*. **a** - FLC protein family. **b -** SVP protein family. The species used in the phylogenetic analysis were: *Quercus suber* (Qs), *Arabidopsis thaliana* (At), *Populus trichocarpa* (Pt), *Quercus robur* (Qr), *Vitis vinifera* (Vv), *Malus x domestica* (Md), *Actinidea sitchensis* (Ac), *Oryza sativa* (Os), *Prunus persica* (Pp), *Beta vulgaris* (Bv), *Arabis alpina* (Aa), *Hordeum vulgare* (Hv), *Medicago truncatula* (Mt), *Cardamine flexuosa* (Cf) and *Glycine max* (Gm). The phylogenies were inferred using the Maximum-likelihood method. The percentage of replicate trees in which the associated taxa clustered together in the bootstrap test (1000 replicates) is shown next to the branches. The evolutionary distances were computed using the JTT matrix-based method and are in the units of the number of amino acid substitutions *per site*. Protein accession numbers used in the phylogenetic analysis are presented in table S1.

In *A. thaliana*, there is only one *SVP* gene but in several other angiosperm species there are several duplicates of *SVP*-like genes (*e.g. P. persica*, *A. chinensis* or *V. vinifera*). Blasting the sequence of SVP against the cork oak database four SVP-like proteins (QsSVP1*-*4) were retrieved and used in the phylogenetic analysis. The SVP sub-family can be divided into two major groups, one containing SVP-like proteins and the other containing DORMANCY ASSOCIATED MADS (DAM) – like proteins. All the QsSVP were included in the SVP-like protein clade, with QsSVP4 being the closest homolog to the Arabidopsis SVP (Fig. 3b).

### The function of some *Q. suber* flowering time genes is conserved in *A. thaliana*

To assess whether the *Q. suber* flowering time genes function in flowering regulation is conserved, *A. thaliana* plants overexpressing *QsFT, QsSPL4, QsSPL9, QsSPL13, QsSOC1, QsFLC, QsSVP1, QsSVP3* and *QsSVP4* were obtained and flowering time was scored in long-day conditions (n=20 per line). Wild-type plants (WT, Col-0) flowered with 9.7 ± 1.3 leaves (Fig. 4a). Arabidopsis plants overexpressing *QsFT* flowered significantly earlier than WT (4.4 ± 1.1 leaves). A similar early flowering phenotype was observed in plants overexpressing *QsSPL4* (5.9 ± 1.5 leaves). The overexpression of *QsSOC1*, *QsSPL9* and *QsSPL13* did not significantly alter flowering time (Fig. 4a).

**Figure 4.**
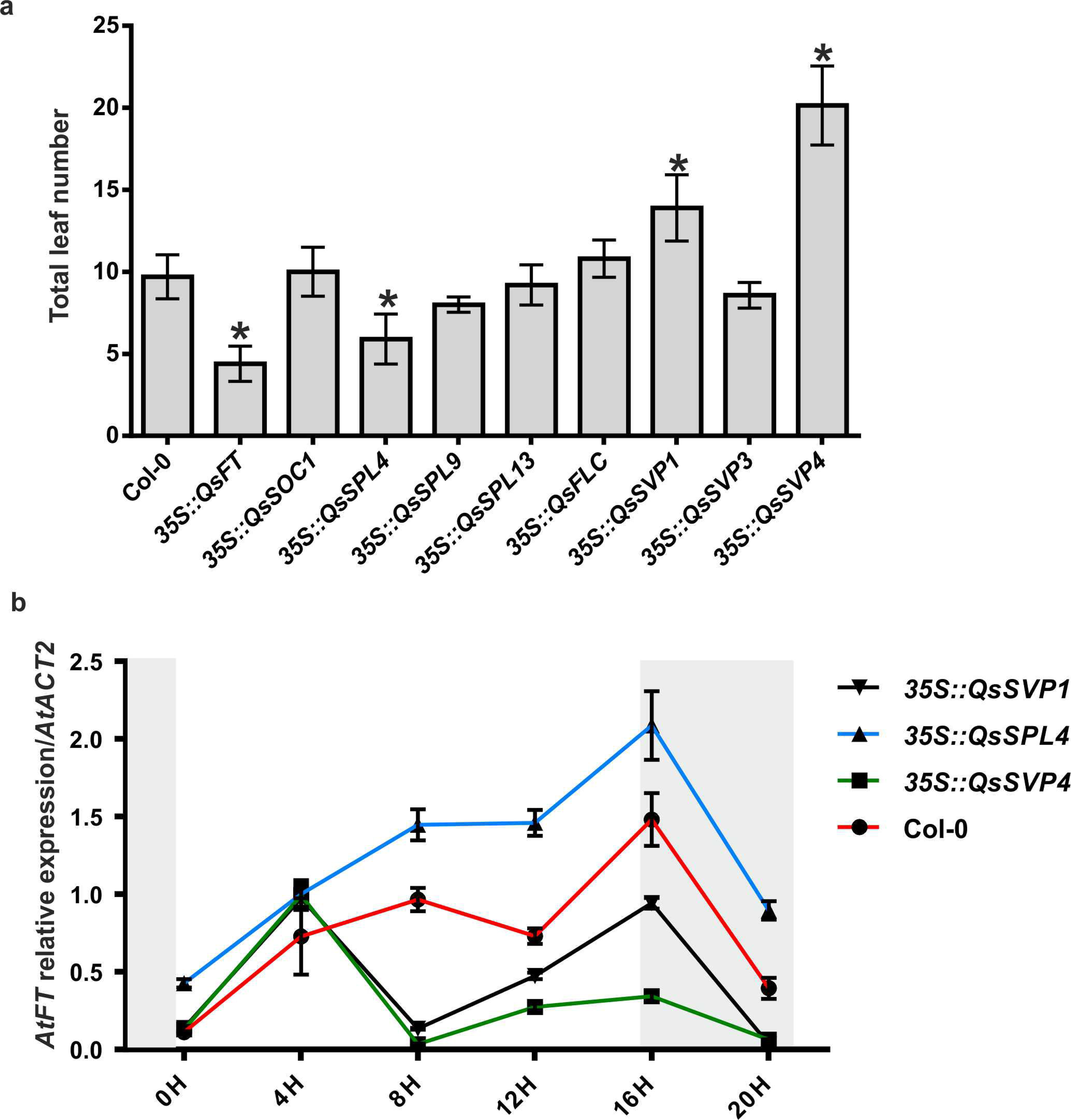
*Quercus suber* flower regulators are able to delay and promote flowering in *Arabidopsis thaliana*. **a** - Total number of leaves of *A. thaliana* wild-type plants (Col-0) and plants overexpressing *QsFT, QsSOC1, QsSPL4, QsSPL9, QsSPL13, QsFLC, QsSVP1, QsSVP3* and *QsSVP4*. Error bars indicate standard deviation (s.d.) of the total number of leaves of 20 plants per independent line. Asterisks indicate p-values ≤ 0.05, determined after performing a Student’s t-test. **b** - *AtFT* expression throughout a 24 hour-course experiment in 10 days-old plants grown under long days and evaluated by qRT-PCR. Col-0 (circle), *35S::SVP1* (inverted triangle); *35S::SPL4* (triangle) and *35S::SVP4* (square). Error bars indicate standard deviation (s.d.) of three biological replicates. *AtACT2* was used as reference gene. White and grey areas indicate day and night periods, respectively.

Plants overexpressing two putative *Q. suber* repressors (*QsFLC* and *QsSVP3*) did not flower later than WT (10.8 ± 1.1 leaves; and 8.6 ± 0.8 leaves, respectively), contrary to the overexpression of *QsSVP1* that generated late-flowering plants (13.9 ± 2.0 leaves) (Fig. 4a). Plants overexpressing *QsSVP4* had a delay in flowering (20.1 +/− 2.4 leaves) (Fig. 4a) and had several morphological defects including extra petals and sepals (Fig. S1b and 1c), suggesting a putative role for *QsSVP4* in flowering time repression and in flower organ patterning.

As a hub for distinct flowering pathways (light/photoperiod, age and vernalization/temperature pathways), the expression profile of *FT* depends on the day length and peaks at the first hours of dark after 16 hours of light (Suárez-López et al. 2001). To study the effect of *Q. suber* flowering regulators in the Arabidopsis *FT* expression, samples of 10-day-old plants of WT and overexpressing lines (that showed altered flowering time, Fig. 4a) were collected in 4-hour intervals during the course of a day (24 hours). WT plants show the typical circadian variation of *FT* expression that peaks during the first hours of dark (Fig. 4b, Col-0). *A. thaliana* plants overexpressing *QsSPL4* have a higher expression of *FT* during the entire course of the experiment suggesting that the early flowering phenotype might derive from the up-regulation of *FT* expression (Fig. 4b). On the contrary, the late flowering phenotypes of the *QsSVP1* and *QsSVP4* overexpressing lines might be correlated with a significant decrease in *FT* expression that culminate in a significantly decreased peak during the first hours of dark (Fig. 4b, *QsSVP1* and *QsSVP4*), which is more evident in the *QsSVP4* overexpressing line.

The results using transgenic *A. thaliana* plants suggest that both *QsFT* and *QsSPL4* are able to induce flowering earlier, whereas *QsSVP1* and *QsSVP4* are able to significantly repress flower induction.

### *Q. suber* flowering-time genes expression fluctuate throughout dormancy and during the growing season

To analyse the expression profile of flowering-time regulators in *Q. suber*, RNA was extracted from leaves and buds of adult and juvenile trees that were chosen according to phenological observations and the levels of expression of the miR156-SPL-miR172 module (Wang et al. 2011, Wang 2014). In several annual and perennial species, the miR156-SPL-miR172 module has been widely used as a marker to establish the plant competence to flower. In cork oak, the levels of miR156 were high in the juvenile and undetected in the adult trees, whereas miR172 displayed the opposite expression pattern (Fig. S2). The expression of both *QsSPL4* and *QsSPL9* in young and adult trees was very similar to miR172 (Fig. S2). This result suggests a conservation of the age module controlled by miR156-SPL-miR172 and reassures that the adults and juvenile trees chosen for sampling can be used to evaluate the trees competence to flower.

To investigate how the putative *Q. suber* flowering-time regulators are involved in the annual flowering cycle, their expression was analysed from growth cessation (September) to bud swelling (March) during two consecutive years. To note, the male flowers adjacent to the swollen axillary buds (March) (Fig. 1d and 1e) were not included in this analysis.

*QsSVP1* and *QsSVP4* gene expression is high in September during axillary bud growth cessation and then steadily decreases until March which is coincident with axillary bud swelling in both the adult and juvenile trees (Fig. 5a and 5b), a pattern that was reproducible in the following year of sampling (Fig. S3a and S3b). *QsSVP3* expression follows a similar pattern to *QsSVP1,* although with a smaller difference between the levels of expression in September and March (Fig. 5a and 5b)*. QsSVP2* expression was not detected during this analysis and *QsFLC* expression did not vary significantly in the sampling period. These results suggest that, similarly to other perennial tree species that undergo dormancy during winter, the *Q. suber SVP*-like genes may be important to the establishment and control of bud growth cessation and burst.

**Figure 5.**
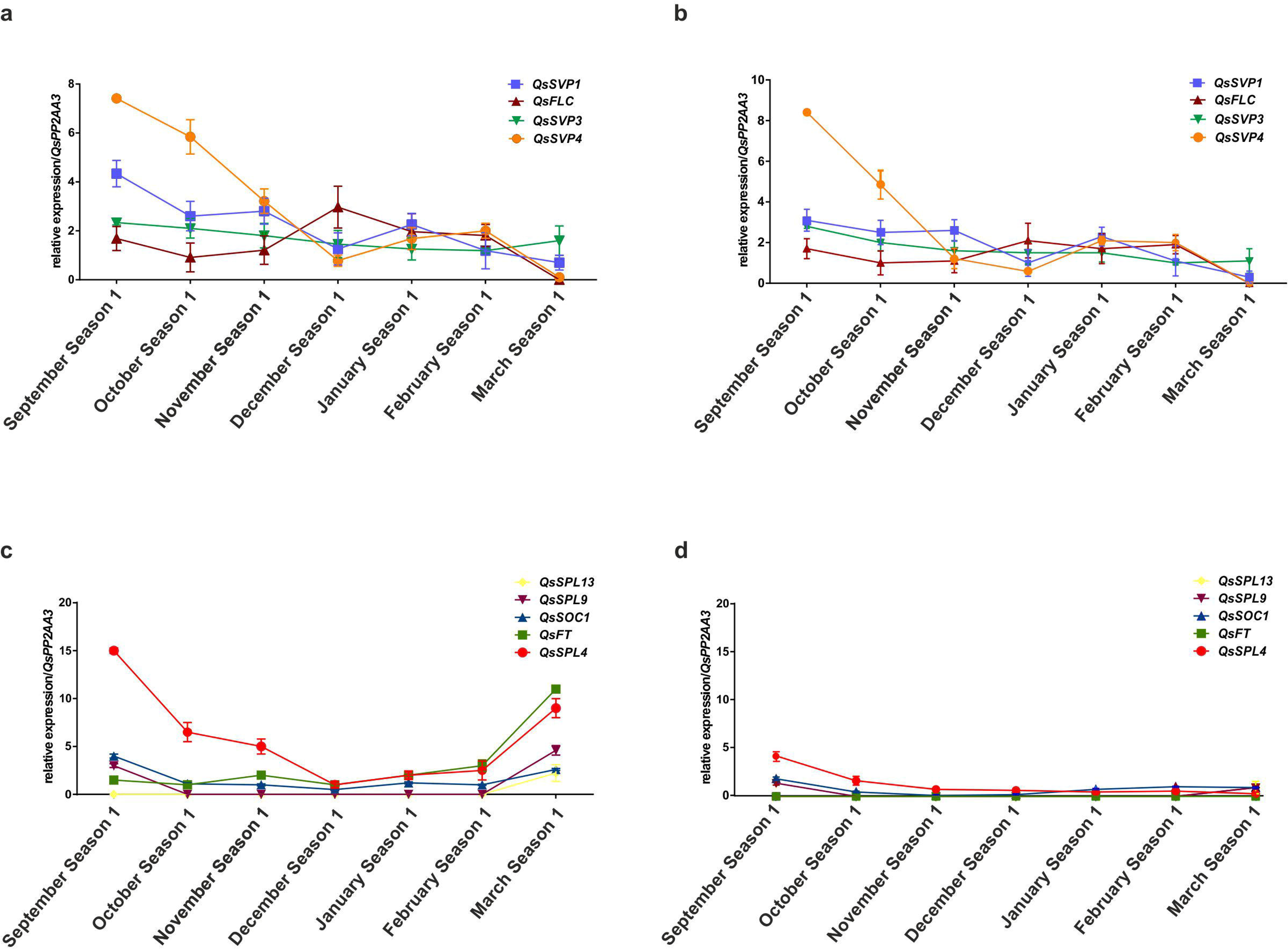
*Quercus suber* flowering-time genes expression in axillary buds from growth cessation to bud swelling. qRT-PCR of *Q. suber* flowering time genes using axillary buds either of adult (a and c) or juvenile trees (b and d), from growth cessation (September) to bud swelling (March). **a** - *QsSVP1*, *QsSVP3*, *QsSVP4* and *QsFLC* expression in axillary buds of adult trees from growth cessation (September) to bud swelling (March). **b -** *QsSVP1*, *QsSVP3*, *QsSVP4* and *QsFLC* expression in axillary buds of the juvenile tree from growth cessation (September) to bud swelling (March). **c** - *QsFT, QsSPL13, QsSPL9*, *QsSOC1* and *QsSPL4* expression in axillary buds of adult trees from growth cessation (September) to bud swelling (March). **d** - *QsFT, QsSPL13, QsSPL9*, *QsSOC1* and *QsSPL4* expression in axillary buds of the juvenile tree from growth cessation (September) to bud swelling (March). Error bars indicate standard deviation (s.d.) of three adult trees and one juvenile tree (three technical replicates per tree). *QsPP2AA3* was used as the reference gene. *QsSVP1* (blue square), *QsSVP3* (green inverted triangle), *QsSVP4* (orange circle)*, QsFLC* (brown triangle), *QsFT* (green square)*, QsSPL13* (yellow square)*, QsSPL9* (brown inverted triangle), *QsSOC1* (purple triangle) and *QsSPL4* (red circle).

The expression of putative flower induction genes was also analysed, both in buds and leaves. The expression of *QsSOC1* did not vary from September (growth cessation) to March (bud swelling) on both adult and juvenile trees (Fig. 5c and Fig. 5d). *QsSPL9* and *QsSPL13* expression was detected in March but only in adult trees (Fig. 5c and Fig. 5d). In adult trees, both *QsFT* and *QsSPL4* expression is high in March during bud swelling (Fig. 5c and Fig. S3c). However, *QsFT* and *QsSPL4* expression was not detected in juveniles in the same period (Fig. 5d and Fig. S3d), thus suggesting that an induction event might be occurring in March during bud swelling.

To investigate further induction events, *QsFT* and *QsSPL4* level of expression was also measured in leaves throughout the year. The expression levels of *QsSPL4* in adult trees leaves did not vary (Fig. 6a). *QsFT* expression varied during the year and had two peaks of expression, one in March and another in August, an expression pattern that was also observed in the following reproductive year (Fig. 6a and Fig. S3e). In juvenile leaves there was no *QsFT* (Fig. 6b and Fig. S3e), suggesting that *QsFT* is likely involved in reproductive induction events, as peaks of expression are observed in adult trees but not in juveniles. The results also suggest that two flowering induction events might be occurring, one in March at the time of bud swelling and another in August before growth cessation. If *QsFT* is behaving as a flower activator it is likely that the two observed peaks of expression may represent the establishment of distinct flower meristems that are temporally separated during the same annual cycle. To assess whether flower meristems are being formed at different times during the year, the expression of the flower meristem identity gene *LFY* was evaluated. *QsLEAFY* was identified in the *Q. suber* transcript database (www.corkoakdb.org) and classified by phylogenetic inference (Fig. S4). *QsLEAFY* expression was only detected in the buds of adult trees, with a higher level of expression in September and in March (Fig. 7a) suggesting that the increased expression of *QsLEAFY* is likely related to the emergence of flower meristems in adult trees, and that it may happen at two different times during the year.

**Figure 6.**
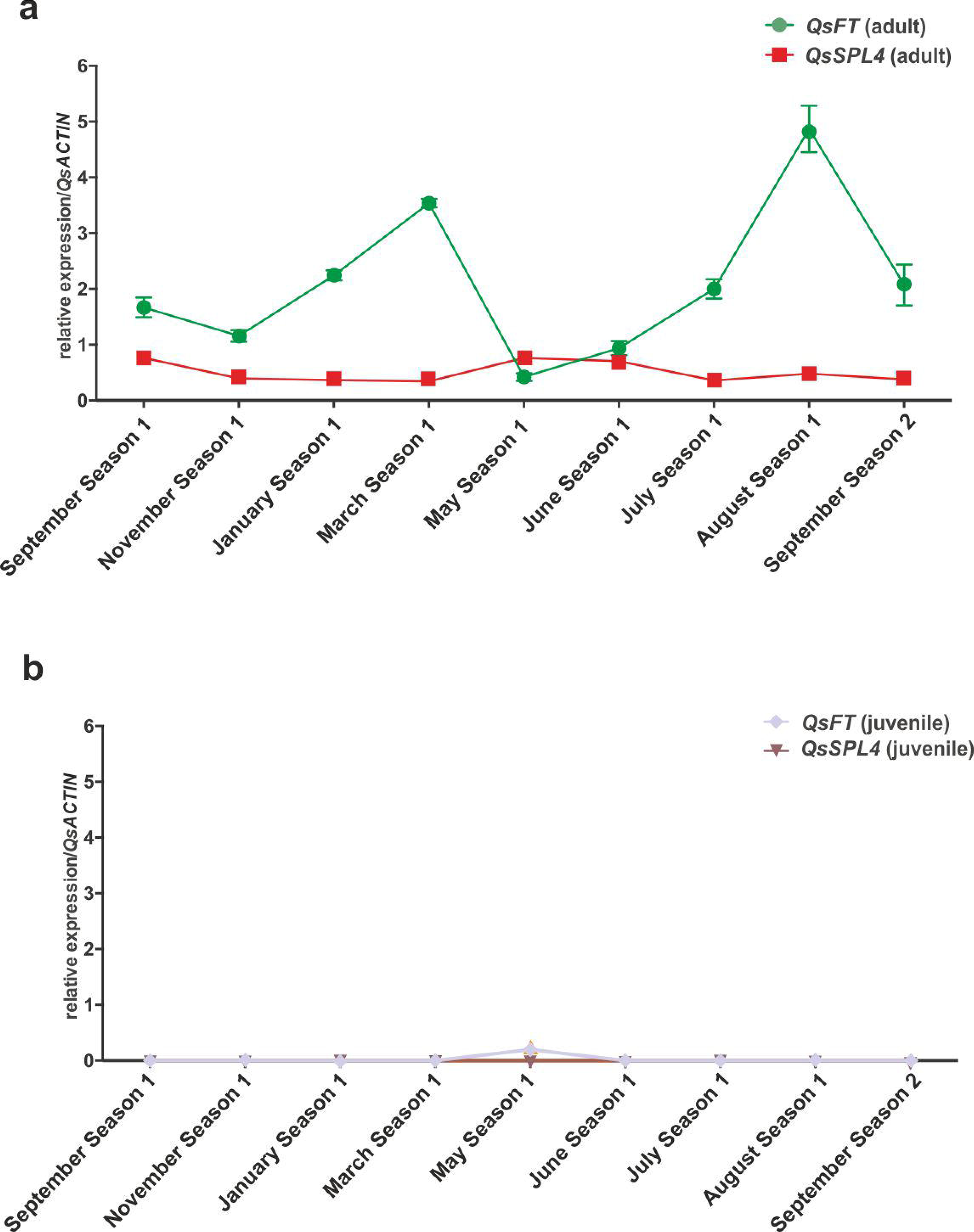
*QsFT* is exclusively expressed in leaves of adult trees. **a** - qRT-PCR of *Q. suber* flowering time genes (*QsFT*, *QsSPL4*) in leaves of adult trees throughout the year*: QsFT* (green circle) and *QsSPL4* (red square). **b** - qRT-PCR of *Q. suber* flowering time genes (*QsFT*, *QsSPL4*) in leaves of a juvenile tree throughout the year*: QsFT* (purple square) and *QsSPL4* (inverted brown triangle). Error bars indicate standard deviation (s.d.) of three adult trees and one juvenile tree (three technical replicates per tree). *QsACTIN* was used as reference gene.

**Figure 7.**
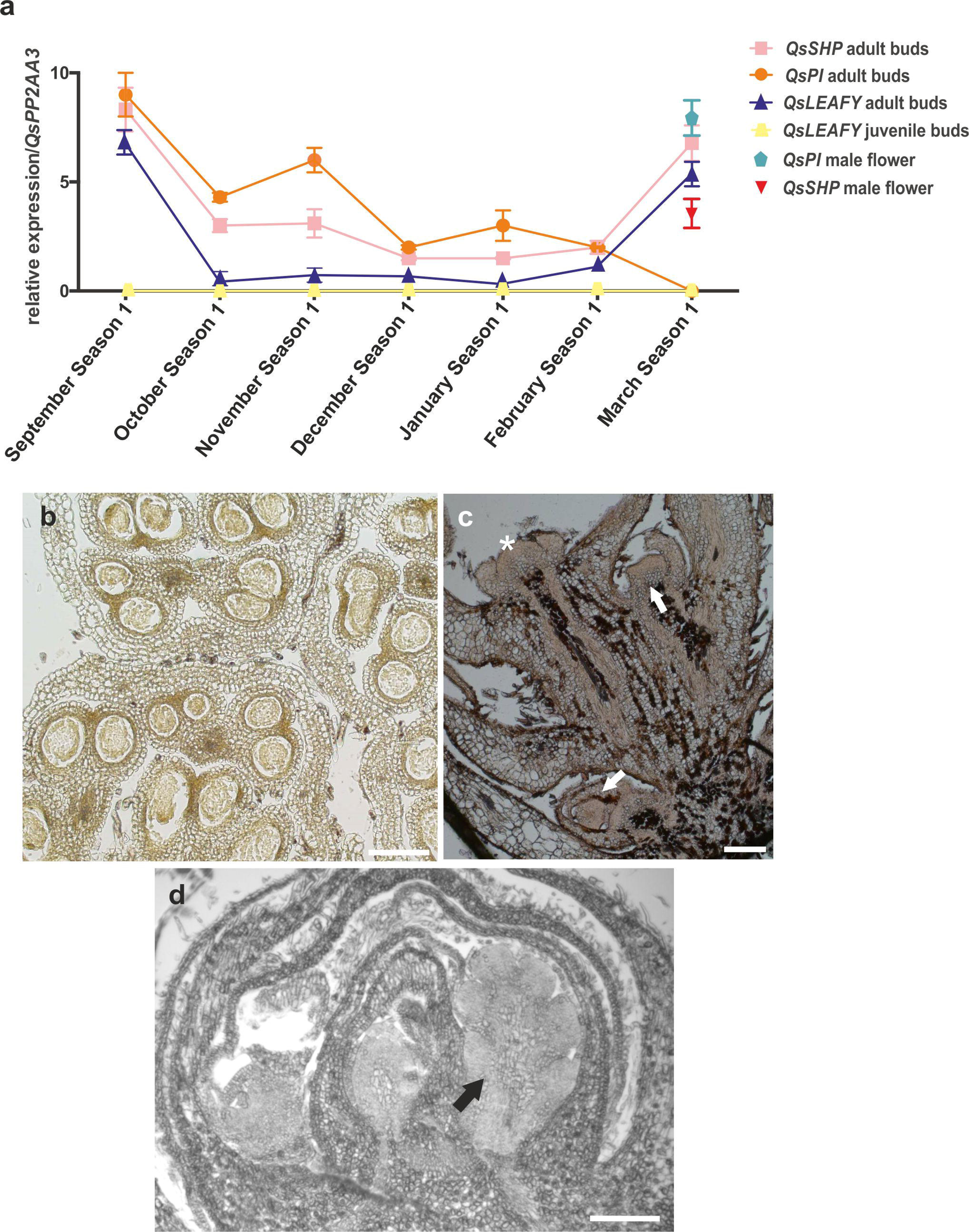
Histological and molecular characterization during bud development. **a** - qRT-PCR of *Q. suber* flower meristem identity (*QsLEAFY*) and flower organ identity genes (*QsPI* and *QsSHP*) of axillary buds collected from September (growth cessation) to March (bud swelling). *QsPI* in axillary buds of adult trees and male flowers (pink circle and blue pentagon, respectively); *QsSHP* in axillary buds of adult trees and male flowers (orange square and red inverted triangle, respectively); and *QsLEAFY* in axillary buds of adult and juvenile trees (dark blue triangle and yellow pentagon, respectively). Error bars indicate standard deviation (s.d.) of three adult trees and one juvenile tree (three technical replicates per tree). *QsPP2AA3* was used as the reference gene. **b -** Cross section of a male flower, that emerged from the swelling axillary bud in March, showing developed anthers. **c -** Longitudinal section of a swelling bud in March containing a shoot axillary meristem (asterisk) and flower meristems (black arrows) in the axils of leaf primordia (black arrows). **d -** Longitudinal section of an axillary bud in September that developed in the axils of new leaves. Inside the axillary bud are inflorescences (black arrow). Scale: 200 μM.

### *Q. suber* male and female flower meristems may be established at different periods

Molecular evidence suggesting the initiation of flower meristems in distinct periods prompted a closer look to the internal organization of the axillary buds in March and September. In the axillary buds of March, the male flower is positioned in the axil of the swollen bud (Fig. 1d, white arrow). The male flower has well defined anthers suggesting an advanced stage of flower development (Fig. 7b). Inside the swelling bud, flower meristems can be found at the axils of leaf primordia (Fig. 7c white arrows). These young flower meristems are likely derived from the flower induction event that occurs in March.

The internal structure of axillary buds in September was also analysed. A section of the axillary bud shows the central meristem protected by scales (Fig. 7d). Outside these first layers of protective scales, are male flower primordia within inflorescences, which are also protected by scales (Fig. 7d, black arrow). Evidence of young inflorescences in September suggests a likely correlation with a flower induction event, which is characterized by a high expression of *QsFT* in leaves during August.

A previous study in *Q. suber* identified several genes, whose expression may be used as a marker for male or female organ development (Rocheta et al. 2014, Sobral and Costa 2017), thus enabling us to infer the sex of the developing inflorescence. According to Sobral and Costa (2017), the B-class MADS-box *QsPISTILLATA* gene (*QsPI*), is uniquely expressed in male flowers, therefore, its expression profile was evaluated during the axillary bud development. *QsPI* was expressed in buds of September and decreased in the following months until no expression was detected in the swollen buds of March (Fig. 7a and Fig. S5a). In March, *QsPI* expression was only detected in the male flowers developing outside the swollen axillary bud (Fig. 7a, blue pentagon). According to Sobral and Costa (2017), the C-class MADS-box *QsSHATTERPROOF* (*QsSHP*) gene is associated to the development of male flowers (together with *QsPI*) and also to the development of the female flowers. *QsSHP* was highly expressed in September, decreased during the winter months and increased again in March, both in the swollen buds and in the male flowers (Fig. 7a and Fig. S5a). Absence of *QsPI* and presence of *QsSHP* transcripts suggests the establishment of a female inflorescence inside the swollen axillary bud in March.

The expression of female flower-specific genes associated to female flower development (*QsWOX9*, *QsDOF1*, *QsKINASE1*, *QsS-LOCUS* and *QsCYP*) (Rocheta et al. 2014) was evaluated in axillary buds in September and March. The expression of some of the female-specific genes was detected in March (bud swelling) but not in September (growth cessation) (Fig. S5b). This result suggests that the inflorescence, present in the axillary bud in September, may be exclusively male. The presence of some female-specific genes and absence of *QsPI* suggests that the inflorescence present in March could be exclusively female.

## Discussion

In monoecious species with accentuated dichogamic systems, female and male flowers develop at different times on the same individual. This reproductive strategy is common to several species across the angiosperms, but little is known about the genetic mechanisms controlling flower induction and what determines the temporal gap between the onset of male and female flowers. According to phenological observations, *Q. suber* is categorized as a protandrous species in which male flower emergence precedes that of female flowers by approximately one to two months. To shed some light on the molecular mechanisms that induce flowering in *Q. suber* and to investigate if they are common to male and female flowers, several putative flower inducer and repressor genes were identified, based on sequence similarities with known regulators of flower development in other species.

In perennial species of temperate climates, a subset of meristems needs to remain vegetative to sustain consecutive growth cycles that are interrupted by unfavourable environmental conditions. Axillary bud growth cessation and bud swelling in perennials, like kiwifruit and peach, is controlled by the regulation of the transcription machinery, particularly by *SVP*-like genes that respond to environmental cues (Li et al. 2009, Wu et al. 2012, 2017). Of the four *QsSVP* genes, only *QsSVP1* and *QsSVP4* were able to delay flowering in transgenic Arabidopsis plants. Furthermore, plants overexpressing *QsSVP4* had flowers with extra petals and sepals. The late flowering and the flower organ number phenotypes derived from overexpressing *QsSVP4* were previously described in other studies in which *SVP*-like genes of several perennial species were overexpressed in perennial and annual species (Li et al. 2010, Wu et al. 2012, 2017, Jaudal et al. 2014). In *Q. suber*, both *QsSVP1* and *QsSVP4* are highly expressed during early stages of axillary bud growth cessation, suggesting a conserved role for these genes in growth arrest. Both *QsSVP1* and *QsSVP4* expression decreased throughout bud dormancy to reach its lower expression levels during axillary bud swelling, similar to what have been described for *SVP* homologs in other perennials such as Castanea and Prunus (Li et al. 2009, Leida et al. 2012, Rios et al. 2014).

Three SPL-like proteins were identified in the cork oak database with homology to the Arabidopsis SPL4, SPL9 and SPL13. *SPL4* and *SPL9* genes have been associated with different functions such as age-dependent developmental transitions, flower induction or leaf morphogenesis (Yu et al. 2015). In leaves of the juvenile tree, the levels of miR156 are high and no *QsSPL4* and *QsSPL9* expression was observed. In adult tree leaves, there was a significant decrease in miR156 levels and increased levels of *QsSPL4, QsSPL9* and miR172 expression. This result suggests that, similarly to other annual and perennial species, the MIR156-SPL-MIR172 module is conserved in the cork oak and that it may be controlling the transition to adulthood.

The role of *QsSPLs* during flower induction was tested by analysing Arabidopsis plants overexpressing *QsSPL4*, *QsSPL9* and *QsSPL13*. Overexpressing *QsSPL13* in Arabidopsis did not alter flowering time, a result that was expected because *SPL13-*like genes have not been, to this date, associated to flower induction (Martin et al. 2010). In Arabidopsis, *SPL9* activates the floral meristem identity genes *FUL* and *SOC1* to control flowering time, but Arabidopsis plants overexpressing *SPL9* do not flower earlier than WT (Wang, Czech, et al. 2009, Wu et al. 2009). Likewise, Arabidopsis plants overexpressing *QsSPL9* had the same number of leaves than WT. Thus, it is not possible to associate a role of *QsSPL9* during floral transition in cork oak.

*QsSPL4* overexpression promotes early flowering in Arabidopsis. The early flowering phenotype is likely due to increased levels of *FT* expression, which is consistent with the known role of *SPL3* (a gene situated in the same clade as *QsSPL4*) in *FT* regulation (Wang, Czech, et al. 2009, Yamaguchi et al. 2009). In cork oak adult trees, *QsSPL4* is expressed in leaves but has a significant increased expression during axillary bud growth cessation and axillary bud swelling. In juvenile trees, *QsSPL4* is also expressed during axillary bud growth cessation. The higher *SPL4* expression during growth cessation is also observed in other perennial species as suggested by transcriptomic data obtained from early dormant buds of poplar and leafy spurge (Ruttink et al. 2007, Doğramacı et al. 2013). Expression of *QsSPL4* during axillary bud growth cessation in both adults and juveniles suggests that *QsSPL4* might be important in a genetic mechanism controlling growth arrest in cork oak. However, *QsSPL4* expression was not detected during axillary bud swelling in juvenile cork oaks, thus suggesting that *QsSPL4* may function in the integration of the flowering induction signals and could be involved in a flowering induction event that might be coincident with the cork oak axillary bud swelling.

Several studies conducted in annual and perennial species have shown that *FT*-like genes are able to induce flowering, as well as accelerate the transition from juvenility to maturity or regulate cycles of seasonal growth (Endo et al. 2005, 2018, Böhlenius et al. 2006, Hsu et al. 2006, 2011, Tränkner et al. 2010, Ye et al. 2014, Fu et al. 2014, Freiman et al. 2015, Zhang et al. 2016, Klocko et al. 2016, Voogd et al. 2017, Miyazaki and Satake 2017, Satake, Kawatsu, Teshima, et al. 2019, Satake, Kawatsu, Chiba, et al. 2019). To confirm that a flower induction event is occurring during axillary bud swelling in cork oak the homolog of *FT* was identified. *QsFT* overexpression induces early flowering in *A. thaliana*, which suggests a conserved role in the vegetative to reproductive transition. *QsFT* expression is low in buds and leaves during growth arrest and throughout winter but is high during axillary bud swelling in March (only in adult trees), thus confirming the existence of a flower induction event.

Using leaf samples throughout the year, another increase in *QsFT* expression was also observed before axillary bud growth cessation in adult trees but not in juvenile trees, suggesting that a second flower induction event could be occurring in that period. One might argue that *QsFT* could have a dual role similarly to the poplar *FT* genes in the control of flowering induction and growth cessation (Böhlenius et al. 2006, Hsu et al. 2011). However, and contrary to what happens in other perennials, such as citrus or apple (Nishikawa et al., 2007; Kotoda et al., 2010), only one canonical *FT* was identified in the cork oak transcript database and the cork oak genome. To confirm the existence of active flower meristems derived from higher *QsFT* expression and thus the proposed flower induction events, the expression of cork oak *LEAFY* was obtained using samples collected from bud growth cessation to bud swelling. The expression profile of *QsLFY* in adult trees coincided with the flowering induction events (one during growth cessation and another during bud swelling). As there was no expression of *QsLFY* in juveniles it is likely that flower meristems are present at different periods during the growth season.

Taking into consideration the *Q. suber* dichogamic nature, it is possible that the two flower induction events might represent a complete separation of the male and female flowers development similarly to what has been described in phenological studies in pecan (Shuhart 1927, Woodroof 1927). Sobral and Costa (2017) suggested that unisexuality in *Q. suber* might derive from a rearrangement of the canonical ABCDE model of flower identity. In that sense, male flowers develop if the B-class gene (*QsPI*) and C-class gene (*QsSHP*) are co-expressed in the same flower meristem, whereas female flowers develop only if *QsSHP* is expressed and *QsPI* is absent. Thus, the *Q. suber* flower meristem derived from the August flowering induction event originates male flowers due to the expression of both *QsPI* and *QsSHP* within the axillary bud. Female inflorescences do not develop during axillary bud growth cessation as suggested by the lack of a carpel-like organ and by the absence of female-specific gene expression. During the March flower induction event that coincides with axillary bud swelling, the absence of *QsPI* expression in the swollen axillary buds may prevent the development of male flowers and enable the progress of the carpel developmental program (Fig. 8). If this is indeed the case, the control of *QsPI* expression after both induction events assumes a pivotal role in the reproductive habit of cork oak.

**Figure 8.**
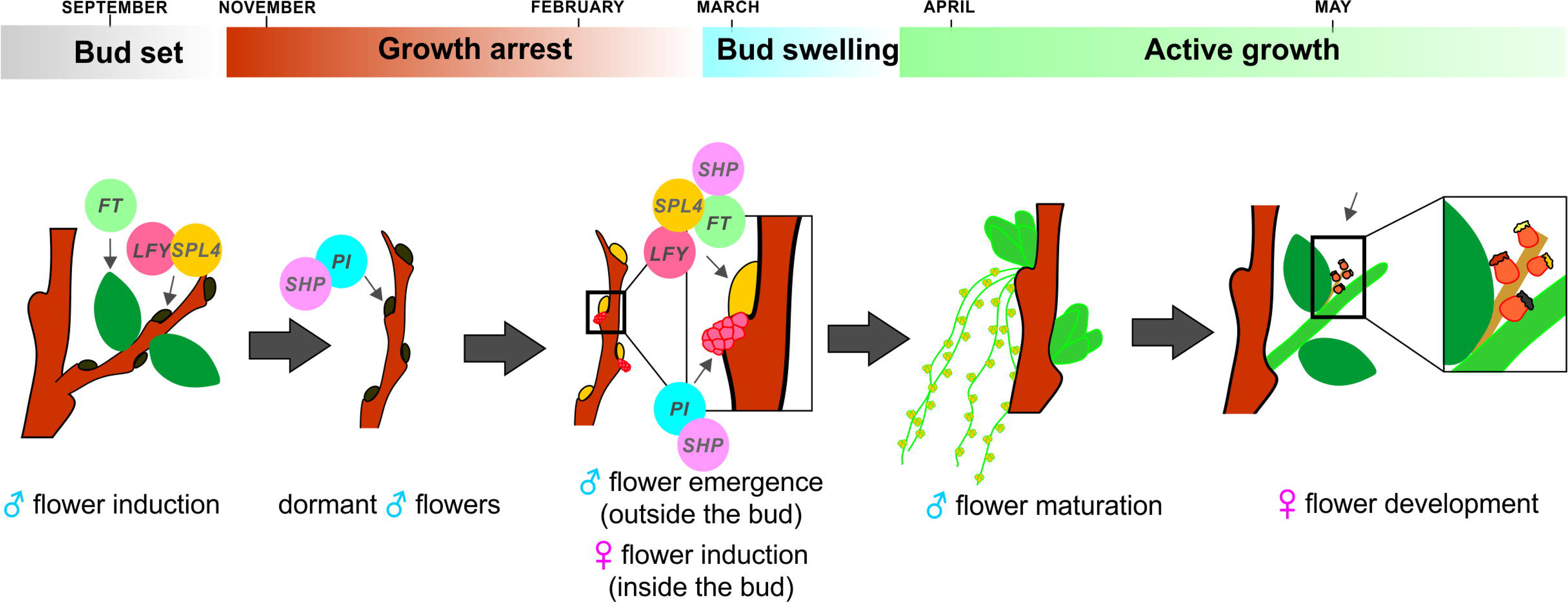
Model for *Quercus suber* flower initiation. The *Q. suber* heterodichogamic habit is characterized by a reproductive season temporally separated in distinct phases. A flower meristem develops (higher *QsLFY* expression) from an induction event in August (higher *QsFT* expression) followed by the up-regulation of growth arrest mechanisms (growth cessation). The inflorescences are exclusively male as *QsPI* is expressed. The male inflorescences remain enclosed in the axillary bud during the growth arrest period. Male flowers will fully develop in March after bud swelling and bud swelling and at the same time of the new vegetative flush. A new flowering event (second peak of *QsFT* expression) occurs during the development of the new leaves that will lead to the induction of female flowers (no *QsPI* expression).

No genetic mechanism has yet been proposed to explain the development of separate flowers that derived from distinct flowering induction events. Therefore, the genetic data gathered here might be useful to explain the heterodichogamic habits common to other woody perennial species.

## Acknowledgements

This work was funded by FCT /COMPETE / FEDER with the project grants FCOMP - 01 - 0124 - FEDER - 019461 / PTDC / AGR-GPL / 118508 / 2010. “Characterization of Reproductive Development of *Quercus suber*”. R.S. was supported by funding from FCT with a Ph.D. grant (ref. SFRH/BD/84365/2012). H.S. was supported by funding from FCT with a Ph.D. grant (ref. SFRH/BD/111529/2015). M.M.R.C. was supported by funding from FCT with a grant (ref. SFRH/BSAB/113781/2015).

## Author contributions

R.S. designed and performed most of the experiments, M.M.R.C. and S.L. designed and supervised the experiments; H.S. performed the the morphological and histological characterization;. R.S. wrote the article with contributions from all the authors.

---

**Figure S1 – *QsSVP4* overexpression in *Arabidopsis thaliana* generates flowers with extra sepals and petals.** a - Col-0 flower. b - Top view of a *35S::QsSVP4* flower. C - Bottom view of *35S::QsSVP4* flower. Scale: 1 mm.

**Figure S2** - ***Quercus suber* flower regulators control the juvenile to adult transition in *Q. suber*.** Gene expression analysis by qRT-PCR of *miR156*, *miR172*, *QsSPL9* and *QsSPL4* in leaves of juvenile and adult trees. Error bars indicate standard deviation (s.d.) of three adult and one juvenile trees (three technical replicates per tree). *QsACTIN* was used as reference gene.

**Figure S3 - Gene expression analysis by qRT-PCR of *Q. suber* flowering time genes** qRT-PCR of *Q. suber* flowering time genes using axillary buds either of adult (a and c) or juvenile trees (b and d) from growth cessation (September) to bud swelling (March). **a** - *QsSVP1*, *QsSVP3*, *QsSVP4* and *QsFLC* expression in axillary buds of adult trees from growth cessation (September) to bud swelling (March). **b -** *QsSVP1*, *QsSVP3*, *QsSVP4* and *QsFLC* expression in axillary buds of the juvenile tree from growth cessation (September) to bud swelling (March). **c** - *QsFT, QsSPL13, QsSPL9*, *QsSOC1* and *QsSPL4* expression in axillary buds of adult trees from growth cessation (September) to bud swelling (March). **d** - *QsFT, QsSPL13, QsSPL9*, *QsSOC1* and *QsSPL4* expression in axillary buds of the juvenile tree from growth cessation (September) to bud swelling (March). Error bars indicate standard deviation (s.d.) of three adult trees and one juvenile tree (three technical replicates per tree). *QsPP2AA3* was used as the reference gene. *QsSVP1* (blue square), *QsSVP3* (green inverted triangle), *QsSVP4* (orange circle)*, QsFLC* (brown triangle), *QsFT* (green square)*, QsSPL13* (yellow square)*, QsSPL9* (brown inverted triangle), *QsSOC1* (purple triangle) and *QsSPL4* (red circle). **e** - Gene expression analysis by qRT-PCR of *QsFT* in leaves of adult and juvenile trees throughout the year*: QsFT* (adult trees, green circle) and *QsFT* (juvenile tree, purple square). Error bars indicate standard deviation (s.d.) of three adult trees and one juvenile tree (three technical replicates per tree). *QsACTIN* was used as reference gene.

**Figure S4 – Homolog of LEAFY in *Quercus suber*.** The *Q. suber* LEAFY homolog is underlined. The phylogeny was inferred using the Maximum-likelihood method. The percentage of replicate trees in which the associated taxa clustered together in the bootstrap test (1000 replicates) is shown next to the branches. The evolutionary distances were computed using the JTT matrix-based method and are in the units of the number of amino acid substitutions *per site*. The species used in the phylogenetic analysis were: *Quercus suber* (Qs), *Arabidopsis thaliana* (At), *Populus trichocarpa* (Pt), *Quercus robur* (Qr), *Malus x domestica* (Md), *Solanum lycopersicum* (Sl), *Oryza sativa* (Os), *Prunus persica* (Pp), *Fagus crenata* (Fc), *Betula pendula* (Bp), *Castanea mollissima* (Cm), *Medicago truncatula* (Mt), *Antirrhinum majus* (Am), *Citrus unshui* and *Juglans regia* (Jr). Protein accession numbers used in the phylogenetic analysis are presented in table S1.

**Figure S5 – Gene expression of *Quercus suber* flower regulators from growth cessation to bud swelling. a -** Gene expression analysis by qRT-PCR of *Q. suber* flower identity genes (*QsPI* and *QsSHP*) of axillary buds collected from growth cessation to bud release (October to March). *QsPI* (orange square) *QsSHP* (pink triangle). **b** *-* **q**RT-PCR of *Q. suber* female flower specific genes (*QsCYP, QsS- LOCUS, QsWOX9, QsDOF1* and *QsKINASE1*) was measured in female flowers in spring and during growth cessation (September) and bud swelling (March). Error bars indicate standard deviation (s.d.) of three adult and one juvenile trees (three technical replicates per tree). *QsPP2AA3* was used as the reference gene.

## References

Abe M, Kobayashi Y, Yamamoto S, Daimon Y, Yamaguchi A, Ikeda Y, Ichinoki H, Notaguchi M, Goto K, Araki T (2005) FD, a bZIP Protein Mediating Signals from the Floral Pathway Integrator FT at the Shoot Apex. Science 309:1052–1056.

Azeez A, Miskolczi P, Tylewicz S, Bhalerao RP (2014) A tree ortholog of APETALA1 mediates photoperiodic control of seasonal growth. Curr Biol 24:717–724.

Azevedo H, Lino-Neto T, Tavares RM (2003) An improved method for high-quality RNA isolation from needles of adult maritime pine trees. Plant Mol Biol Report 21:333–338.

Bai WN, Zeng YF, Zhang DY (2007) Mating patterns and pollen dispersal in a heterodichogamous tree, Juglans mandshurica (Juglandaceae). New Phytol 176:699–707.

Becker A, Theißen G (2003) The major clades of MADS-box genes and their role in the development and evolution of flowering plants. Mol Phylogenet Evol 29:464–489.

Blázquez M, Soowal LN, Lee I, Weigel D (1997) LEAFY expression and flower initiation in Arabidopsis. Development 124:3835–3844.

Boavida LC, Varela MC, Feijo JA (1999) Sexual reproduction in the cork oak (Quercus suber L.). I. The progamic phase. Sex Plant Reprod 11:347–353.

Böhlenius H, Huang T, Charbonnel-Campaa L, Brunner AM, Jansson S, Strauss SH, Nilsson O (2006) CO/FT regulatory module controls timing of flowering and seasonal growth cessation in trees. Science 312:1040–1043.

Bowman JL, Alvarez J, Weigel D, Meyerowitz EM, Smyth DR (1993) Control of flower development in Arabidopsis thaliana by APETALA1 and interacting genes. Development 119:721–743.

Bowman JL, Smyth DR, Meyerowitz EM (1989) Genes directing flower development in Arabidopsis. Plant Cell 1:37–52.

Capovilla G, Schmid M, Posé D (2015) Control of flowering by ambient temperature. J Exp Bot 66:59–69.

Chang SJ, Puryear J, Cairney J (1993) A simple and efficient method for isolating RNA from pine trees. Plant Mol Biol Report 11:113–116.

Clough SJ, Bent AF (1998) Floral dip: A simplified method for Agrobacterium-mediated transformation of Arabidopsis thaliana. Plant J 16:735–743.

Coen ES, Meyerowitz EM (1991) The war of the whorls: genetic interactions controlling flower development. Nature 353:31–37.

Derory J, Léger P, Garcia V, Schaeffer J, Hauser MT, Salin F, Luschnig C, Plomion C, Glössl J, Kremer A (2006) Transcriptome analysis of bud burst in sessile oak (Quercus petraea). New Phytol 170:723–738.

Doğramacı M, Foley ME, Chao WS, Christoffers MJ, Anderson J V (2013) Induction of endodormancy in crown buds of leafy spurge (Euphorbia esula L.) implicates a role for ethylene and cross-talk between photoperiod and temperature. Plant Mol Biol 81:577– 593.

Elena-Rossello JA, de Rio JM, Valdecantos Garcia JL, Santamaria IG (1993) Ecological aspects of the floral phenology of the cork-oak (Q suber L): why do annual and biennial biotypes appear?

Endo T, Shimada T, Fujii H, Kobayashi Y, Araki T, Omura M (2005) Ectopic expression of an FT homolog from Citrus confers an early flowering phenotype on trifoliate orange (Poncirus trifoliata L. Raf.). Transgenic Res 14:703–712.

Endo T, Shimada T, Nakata Y, Fujii H, Matsumoto H, Nakajima N, Ikoma Y, Omura M (2018) Abscisic acid affects expression of citrus FT homologs upon floral induction by low temperature in Satsuma Mandarin (Citrus unshiu Marc.). Tree Physiol 38:755–771.

Felsenstein J (1986) PHYLIP -- Phylogeny Inference Package (Version 3.2). Cladistics

Flanagan C, Hu Y, Map H (1996) Specific expression of the AGL1 MADS-box gene suggests regulatory functions in Arabidopsis gynoecium and ovule development. Plant J 10:343–353.

Freiman A, Golobovitch S, Yablovitz Z, Belausov E, Dahan Y, Peer R, Avraham L, Freiman Z, Evenor D, Reuveni M, Sobolev V, Edelman M, Shahak Y, Samach A, Flaishman MA (2015) Expression of flowering locus T2 transgene from Pyrus communis L. delays dormancy and leaf senescence in Malus ?? domestica Borkh, and causes early flowering in tobacco. Plant Sci 241:164–176.

Fu J, Wang L, Wang Y, Yang L, Yang Y, Dai S (2014) Photoperiodic control of FT-like gene ClFT initiates flowering in Chrysanthemum lavandulifolium. Plant Physiol Biochem 74:230–238.

Gleiser G, Verdú M, Segarra-Moragues JG, González-Martínez SC, Pannell JR (2008) Disassortative mating, sexual specialization, and the evolution of gender dimorphism in heterodichogamous Acer opalus. Evolution (NY) 62:1676–1688.

Golembeski GS, Kinmonth-Schultz HA, Song YH, Imaizumi T (2014) Photoperiodic Flowering Regulation in Arabidopsis thaliana. Adv Bot Res 72:1–28.

Gómez-Casero MT, Galán C, Domínguez-Vilches E (2007) Flowering phenology of Mediterranean Quercus species in different locations (Córdoba, SW Iberian Peninsula). Acta botánica Malacit 32:127–146.

Hartmann U, Höhmann S, Nettesheim K, Wisman E, Saedler H, Huijser P (2000) Molecular cloning of SVP: A negative regulator of the floral transition in Arabidopsis. Plant J 21:351–360.

Horvath DP, Sung S, Kim D, Chao W, Anderson J (2010) Characterization, expression and function of DORMANCY ASSOCIATED MADS-BOX genes from leafy spurge. Plant Mol Biol 73:169–179.

Howe GT, Horvath DP, Dharmawardhana P, Priest HD, Mockler TC, Strauss SH (2015) Extensive Transcriptome Changes During Natural Onset and Release of Vegetative Bud Dormancy in Populus. Front Plant Sci 6:1–28.

Hsu C-Y, Adams JP, Kim H, No K, Ma C, Strauss SH, Drnevich J, Vandervelde L, Ellis JD, Rice BM, Wickett N, Gunter LE, Tuskan GA, Brunner AM, Page GP, Barakat A, Carlson JE, DePamphilis CW, Luthe DS, Yuceer C (2011) FLOWERING LOCUS T duplication coordinates reproductive and vegetative growth in perennial poplar. Proc Natl Acad Sci U S A 108:10756–10761.

Hsu CY, Liu Y, Luthe DS, Yuceer C (2006) Poplar *FT2* Shortens the Juvenile Phase and Promotes Seasonal Flowering. Plant Cell 18:1846–1861.

Hwan Lee J, Ryu H-S, Sook Chung K, Posé D, Kim S, Schmid M, Ahn JH (2013) Regulation of Temperature-Responsive Flowering by MADS-Box Transcription Factor Repressors. Science 342:628–632.

Ibáñez C, Kozarewa I, Johansson M, Ogren E, Rohde A, Eriksson ME (2010) Circadian clock components regulate entry and affect exit of seasonal dormancy as well as winter hardiness in Populus trees. Plant Physiol 153:1823–1833.

Jaudal M, Monash J, Zhang L, Wen J, Mysore KS, Macknight R, Putterill J (2014) Overexpression of Medicago SVP genes causes floral defects and delayed flowering in Arabidopsis but only affects floral development in Medicago. J Exp Bot 65:429–442.

Jiménez S, Li Z, Reighard GL, Bielenberg DG (2010) Identification of genes associated with growth cessation and bud dormancy entrance using a dormancy-incapable tree mutant. BMC Plant Biol 10:25–34

Johansson M, Staiger D (2015) Time to flower: Interplay between photoperiod and the circadian clock. J Exp Bot 66:719–730.

Karlberg A, Bako L, Bhalerao RP (2011) Short day-mediated cessation of growth requires the downregulation of AINTEGUMENTALIKE1 transcription factor in hybrid aspen. PLoS Genet 7

Karlgren A, Gyllenstrand N, Clapham D, Lagercrantz U (2013) FLOWERING LOCUS T/TERMINAL FLOWER1-like genes affect growth rhythm and bud set in Norway spruce. Plant Physiol 163:792–803.

Kaul RB (1985) Reproductive Morphology of Quercus (Fagaceae). Am J Bot 72:1962.

Klocko AL, Ma C, Robertson S, Esfandiari E, Nilsson O, Strauss SH (2016) FT overexpression induces precocious flowering and normal reproductive development in Eucalyptus. Plant Biotechnol J 14:808–819.

Kobayashi Y, Kaya H, Goto K, Iwabuchi M, Araki T (1999) A pair of related genes with antagonistic roles in mediating flowering signals. Science 286:1960–1962.

Kotoda N, Hayashi H, Suzuki M, Igarashi M, Hatsuyama Y, Kidou SI, Igasaki T, Nishiguchi M, Yano K, Shimizu T, Takahashi S, Iwanami H, Moriya S, Abe K (2010) Molecular characterization of flowering LOCUS t-like genes of apple (malus X domestica borkh.). Plant Cell Physiol 51:561–575.

Ledig FT, Beland JW, Fryer JH (1971) Breeding techniques for white oak. In: Southern Conf Forest Tree Impr Proc.

Leida C, Conesa A, Llácer G, Badenes ML, Ríos G (2012) Histone modifications and expression of DAM6 gene in peach are modulated during bud dormancy release in a cultivar-dependent manner. New Phytol 193:67–80.

Li D, Liu C, Shen L, Wu Y, Chen H, Robertson M, Helliwell CA, Ito T, Meyerowitz E, Yu H (2008) A Repressor Complex Governs the Integration of Flowering Signals in Arabidopsis. Dev Cell 15:110–120.

Li Z, Reighard GL, Abbott AG, Bielenberg DG (2009) Dormancy-associated MADS genes from the EVG locus of peach [Prunus persica (L.) Batsch] have distinct seasonal and photoperiodic expression patterns. J Exp Bot 60:3521–3530.

Li ZM, Zhang JZ, Mei L, Deng XX, Hu CG, Yao JL (2010) PtSVP, an SVP homolog from trifoliate orange (Poncirus trifoliata L. Raf.), shows seasonal periodicity of meristem determination and affects flower development in transgenic Arabidopsis and tobacco plants. Plant Mol Biol 74:129–142.

Liljegren SJ, Ditta GS, Eshed Y, Savidge B, Bowman JL, Yanofsky MF (2000) SHATTERPROOF MADS-box genes control seed dispersal in Arabidopsis. Nature 404:766–770.

Livak KJ, Schmittgen TD (2001) Analysis of relative gene expression data using real-time quantitative PCR and. Methods 25:402–408.

Löytynoja A, Goldman N (2005) An algorithm for progressive multiple alignment of sequences with insertions. Proc Natl Acad Sci U S A 102:10557–10562.

Martin RC, Asahina M, Liu P-P, Kristof JR, Coppersmith JL, Pluskota WE, Bassel GW, Goloviznina NA, Nguyen TT, Martínez-Andújar C, Arun Kumar MB, Pupel P, Nonogaki H (2010) The regulation of post-germinative transition from the cotyledon- to vegetative-leaf stages by microRNA-targeted SQUAMOSA PROMOTER-BINDING PROTEIN LIKE13 in Arabidopsis. Seed Sci Res 20:89.

Marum L, Miguel A, Ricardo CP, Miguel C (2012) Reference gene selection for quantitative real-time PCR normalization in quercus suber. PLoS One 7

Michaels SD, Amasino RM (1999) FLOWERING LOCUS C encodes a novel MADS domain protein that acts as a repressor of flowering. Plant Cell 11:949–56.

Miyazaki Y, Satake A (2017) Relationship between seasonal progression of floral meristem development and FLOWERING LOCUS T expression in the deciduous tree Fagus crenata. Ecol Res 32:627–631.

Moyroud E, Minguet EG, Ott F, Yant L, Posé D, Monniaux M, Blanchet S, Bastien O, Thévenon E, Weigel D, Schmid M, Parcy FF (2011) Prediction of regulatory interactions from genome sequences using a biophysical model for the Arabidopsis LEAFY transcription factor. Plant Cell 23:1293–306.

Natividade J (1950) Subericultura (In portuguese).

Neale DB, Kremer A (2011) Forest tree genomics: growing resources and applications. Nat Rev Genet 12:111–122.

Nishikawa F, Endo T, Shimada T, Fujii H, Shimizu T, Omura M (2007) Increased CiFT abundance in the stem correlates with floral induction by low temperature in Satsuma mandarin (Citrus unshiu Marc.). 58:3915–3927.

Oliveira G, Correia O, Martins-Loução MA, Catarino FM (1994) Phenological and growth patterns of the Mediterranean oak Quercus suber L. Trees 9:41–46.

Parcy F, Nilsson O, Busch MA, Lee I, Weigel D (1998) A genetic framework for floral patterning. Nature 395:561–566.

Pelaz S, Ditta GS, Baumann E, Wisman E, Yanofsky MF (2000) B and C floral organ identity functions require SEPALLATA MADS-box genes. Nature 405:200–203.

Peña L, Martín-Trillo M, Juárez J, Pina J a, Navarro L, Martínez-Zapater JM (2001) Constitutive expression of Arabidopsis LEAFY or APETALA1 genes in citrus reduces their generation time. Nat Biotechnol 19:263–267.

Pendleton RL, Freeman DC, McArthur ED, Sanderson SC (2000) Gender specialization in heterodichogamous Grayia brandegei (Chenopodiaceae): Evidence for an alternative pathway to dioecy. Am J Bot 87:508–516.

Polito VS, Li NY (1985) Pistil late flower differentiation in English walnut (Juglans regia L.): A developmental basis for heterodichogamy. Sci Hortic (Amsterdam) 26:333–338.

Posé D, Verhage L, Ott F, Yant L, Mathieu J, Angenent GC, Immink RGH, Schmid M (2013) Temperature-dependent regulation of flowering by antagonistic FLM variants. Nature 503:414–7.

Renner SS, Beenken L, Grimm GW, Kocyan A, Ricklefs RE (2007) The evolution of dioecy, heterodichogamy, and labile sex expression in Acer. Evolution (N Y) 61:2701–2719.

Rios G, Leida C, Conejero A, Badenes ML (2014) Epigenetic regulation of bud dormancy events in perennial plants. Front Plant Sci 5:247.

Rocheta M, Sobral R, Magalhães J, Amorim MI, Ribeiro T, Pinheiro M, Egas C, Morais-Cecílio L, Costa MMR (2014) Comparative transcriptomic analysis of male and female flowers of monoecious Quercus suber. Front Plant Sci 5:1–16.

Rohde, Howe G, Olsen J, Moritz T, Van Montagu M, Junttila O, Boerjan W (2000) Molecular aspects of bud dormancy in trees. Mol Biol Woody Plants 1:89–134. http://link.springer.com/chapter/10.1007/978-94-017-2311-4_4

Rottmann WH, Meilan R, Sheppard LA, Brunner AM, Skinner JS, Ma C, Cheng S, Jouanin L, Pilate G, Strauss SH (2000) Diverse effects of overexpression of LEAFY and PTLF, a poplar (Populus) homolog of LEAFY/FLORICAULA, in transgenic poplar and Arabidopsis. Plant J 22:235–245.

Ruttink T, Arend M, Morreel K, Storme V, Rombauts S, Fromm J, Bhalerao RP, Boerjan W, Rohde A (2007) A Molecular Timetable for Apical Bud Formation and Dormancy Induction in Poplar. PLANT CELL ONLINE

Saito T, Bai S, Ito A, Sakamoto D, Saito T, Ubi BE, Imai T, Moriguchi T (2013) Expression and genomic structure of the dormancy-associated MADS box genes MADS13 in Japanese pears (Pyrus pyrifolia Nakai) that differ in their chilling requirement for endodormancy release. Tree Physiol 33:654–667.

Samach A, Onouchi H, Gold SE, Ditta GS, Schwarz-Sommer Z, Yanofsky MF, Coupland G, Koornneef M, Alonso-Blanco C, Peeters AJM, Soppe W, Levy YY, Dean C, Samach A, Coupland G, Koornneef M, Hanhart CJ, Veen JH van der, Hanhart CJ, Peeters AJM, Redei GP, Putterill J, Robson F, Lee K, Simon R, Coupland G, Simon R, Igeño MI, Coupland G, Sablowski RWM, Meyerowitz EM, Kobayashi YKH, Goto K, Iwabuchi M, Araki T, Diatchenko L, Lin X, Menzel G, Apel K, Melzer S, Smyth DR, Bowman JL, Meyerowitz EM, Yanofsky MF, Strizhov N, Guzman P, Ecker JR, Nanjo TKM, Nilsson O, Lee I, Blázquez MA, Weigel D, Macknight R, Kardailsky I, Ruiz-Garcia L, Sheldon CC, Michaels SD, Amasino RM, Sheldon CC, Rouse DT, Finnegan EJ, Peacock WJ, Dennis ES, Corbesier L, Gadisseur I, Silvestre G, Jacqmard A, Bernier G, Liljegren SJ, Gustafson-Brown C, Pinyopich A, Ditta GS, Yanofsky MF (2000) Distinct roles of CONSTANS target genes in reproductive development of Arabidopsis. Science 288:1613–616.

Samach A, Smith HM (2013) Constraints to obtaining consistent annual yields in perennials. II: Environment and fruit load affect induction of flowering. Plant Sci 207:168–176.

Satake A, Kawatsu K, Chiba Y, Kitamura K, Han Q (2019) Synchronized expression of FLOWERING LOCUS T between branches underlies mass flowering in Fagus crenata. Popul Ecol 61:5–13.

Satake A, Kawatsu K, Teshima K, Kabeya D, Han Q (2019) Field transcriptome revealed a novel relationship between nitrate transport and flowering in Japanese beech. Sci Rep 9:1–12.

Schwarz S, Grande A V., Bujdoso N, Saedler H, Huijser P (2008) The microRNA regulated SBP-box genes SPL9 and SPL15 control shoot maturation in Arabidopsis. Plant Mol Biol 67:183–195.

Sedgley M, Griffin AR (1989) Sexual Reproduction of Tree Crops. http://www.sciencedirect.com/science/article/pii/B978012634470750012X

Sheldon CC, Rouse DT, Finnegan EJ, Peacock WJ, Dennis ES (2000) The molecular basis of vernalization: the central role of Flowering Locus (FLC). Proc Natl Acad Sci U S A 97:3753–3758.

Shuhart DV (1927) the Morphological Differentiation of the Pistillate Flowers of the Pecan ^. 4

Sobral R, Costa MMR (2017) Role of floral organ identity genes in the development of unisexual flowers of Quercus suber L. Sci Rep 7:10368–10475

Song YH, Shim JS, Kinmonth-Schultz H, Imaizumi T (2015) Photoperiodic Flowering: Time Measurement Mechanisms in Leaves. Annu Rev Plant Biol 66:441–464.

Suárez-López P, Wheatley K, Robson F, Onouchi H, Valverde F, Coupland G (2001) CONSTANS mediates between the circadian clock and the control of flowering in Arabidopsis. Nature 410:1116–1120.

Tao Z, Shen L, Liu C, Liu L, Yan Y, Yu H (2012) Genome-wide identification of SOC1 and SVP targets during the floral transition in Arabidopsis. Plant J 70:549–561.

Taoka K, Ohki I, Tsuji H, Furuita K, Hayashi K, Yanase T, Yamaguchi M, Nakashima C, Purwestri YA, Tamaki S, Ogaki Y, Shimada C, Nakagawa A, Kojima C, Shimamoto K (2011) 14-3-3 proteins act as intracellular receptors for rice Hd3a florigen. Nature 476:332–335.

Tisserat B, Esan EB, Murashige T (1979) Somatic embryogenesis in angiosperms. Hortic Rev (Am Soc Hortic Sci) 1:1–78.

Tränkner C, Lehmann S, Hoenicka H, Hanke MV, Fladung M, Lenhardt D, Dunemann F, Gau A, Schlangen K, Malnoy M, Flachowsky H (2010) Over-expression of an FT-homologous gene of apple induces early xowering in annual and perennial plants. Planta 232:1309–1324.

Tylewicz S, Tsuji H, Miskolczi P, Petterle A, Azeez A, Jonsson K, Shimamoto K, Bhalerao RP (2015) Dual role of tree florigen activation complex component FD in photoperiodic growth control and adaptive response pathways. Proc Natl Acad Sci U S A 112:3140–5.

Varela MC, Valdiviesso T (1996) Phenological phases of Quercus suber L. flowering. For Genet 3:93–102.

Varkonyi-Gasic E, Moss SM, Voogd C, Wu R, Lough RH, Wang Y-Y, Hellens RP (2011) Identification and characterization of flowering genes in kiwifruit: sequence conservation and role in kiwifruit flower development. BMC Plant Biol 11:72.

Voogd C, Brian LA, Wang T, Allan AC, Varkonyi-Gasic E (2017) Three FT and multiple CEN and BFT genes regulate maturity, flowering, and vegetative phenology in kiwifruit. J Exp Bot 68:1539–1553.

Wang JW (2014) Regulation of flowering time by the miR156-mediated age pathway. J Exp Bot 65:4723–4730.

Wang JW, Czech B, Weigel D (2009) miR156-Regulated SPL Transcription Factors Define an Endogenous Flowering Pathway in Arabidopsis thaliana. Cell 138:738–749.

Wang R, Farrona S, Vincent C, Joecker A, Schoof H, Turck F, Alonso-Blanco C, Coupland G, Albani MC (2009) PEP1 regulates perennial flowering in Arabis alpina. Nature 459:423–427.

Wang JW, Park MY, Wang LJ, Koo Y, Chen XY, Weigel D, Poethig RS (2011) MiRNA control of vegetative phase change in trees. PLoS Genet 7

Watanabe S, Noma N, Nishida T (2016) Flowering phenology and mating success of the heterodichogamous tree Machilus thunbergii Sieb. et Zucc (Lauraceae). Plant Species Biol 31:29–37.

Weigel D, Nilsson O (1995) A developmental switch sufficient for flower initiation in diverse plants. Nature 377:495–500.

Wester PJ (1910) Pollination experiments with anonas. Bull Torrey Bot Club 37:529–539.

Wigge PA, Kim MC, Jaeger KE, Busch W, Schmid M, Lohmann JU, Weigel D (2005) Integration of spatial and temporal information during floral induction in Arabidopsis. Science 309:1056–1059.

Winter CM, Austin RS, Blanvillain-Baufumé S, Reback MA, Monniaux M, Wu MF, Sang Y, Yamaguchi A, Yamaguchi N, Parker JE, Parcy F, Jensen ST, Li H, Wagner D (2011) LEAFY Target Genes Reveal Floral Regulatory Logic, cis Motifs, and a Link to Biotic Stimulus Response. Dev Cell 20:430–443.

Woodroof BJG (1927) From flower to maturity ^. 34:1049–1064.

Wu G, Park MY, Conway SR, Wang JW, Weigel D, Poethig RS (2009) The Sequential Action of miR156 and miR172 Regulates Developmental Timing in Arabidopsis. Cell 138:750–759.

Wu G, Poethig RS (2006) Temporal regulation of shoot development in Arabidopsis thaliana by miR156 and its target SPL3. Development 133:3539–3547.

Wu R, Tomes S, Karunairetnam S, Tustin SD, Hellens RP, Allan AC, Macknight RC, Varkonyi-Gasic E (2017) SVP-like MADS Box Genes Control Dormancy and Budbreak in Apple. Front Plant Sci 08.

Wu RM, Walton EF, Richardson AC, Wood M, Hellens RP, Varkonyi-Gasic E (2012) Conservation and divergence of four kiwifruit SVP-like MADS-box genes suggest distinct roles in kiwifruit bud dormancy and flowering. J Exp Bot 63:797–807.

Yamaguchi A, Wu MF, Yang L, Wu G, Poethig RS, Wagner D (2009) The MicroRNA-Regulated SBP-Box Transcription Factor SPL3 Is a Direct Upstream Activator of LEAFY, FRUITFULL, and APETALA1. Dev Cell 17:268–278.

Yamane H, Ooka T, Jotatsu H, Hosaka Y, Sasaki R, Tao R (2011) Expressional regulation of PpDAM5 and PpDAM6, peach (Prunus persica) dormancy-associated MADS-box genes, by low temperature and dormancy-breaking reagent treatment. J Exp Bot 62:3481–3488.

Ye J, Geng Y, Zhang B, Mao H, Qu J, Chua N (2014) The Jatropha FT ortholog is a systemic signal regulating growth and flowering time. Biotechnol Biofuels 7:91.

Yu S, Lian H, Wang JW (2015) Plant developmental transitions: The role of microRNAs and sugars. Curr Opin Plant Biol 27:1–7.

Zhang L, Yu H, Lin S, Gao Y (2016) Molecular Characterization of FT and FD Homologs from Eriobotrya deflexa Nakai forma koshunensis. Front Plant Sci 7:1–10.

Zhou C-M, Zhang T-Q, Wang X, Yu S, Lian H, Tang H, Feng Z-Y, Zozomova-Lihova J, Wang J-W (2013) Molecular Basis of Age-Dependent Vernalization in Cardamine flexuosa. Science (80-) 340:1097–1100.

