## Supplemental tables for "Unisexual flower initiation in the monoecious *Quercus suber* L., a molecular approach"

### Supporting information

#### Supporting Table I – List of gene accessions

| *Cork oak* | Corkoak database | *A. thaliana* | NCBI accession | *Vitis vinifera* | NCBI accession |
|---|---|---|---|---|---|
| *QsFLC* | QSP044604.0 | *AtFLC* | AED91498.1 | *VvSPL1* | XP_002275728.1 |
| *QsSOC1* | QSP149164.0 | *AtSOC1* | AEC10583.1 | *VvSPL2* | XP_002270226.1 |
| *QsSPL4* | QSP140838.0 | *AtSVP* | AEC07320.1 | *VvFLC* | ACZ26524.1 |
| *QsSPL9* | QSP053728.0 | *AtSPL4* | CAB56584.1 | ***B. vulgaris*** | **NCBI accession** |
| *QsSPL13* | QSP014890.0 | *AtSPL9* | CAB56591.1 | *BvFT1* | ADM92607.1 |
| *QsFT* | QSP022516.0 | *AtSPL13* | OAO91371.1 | *BvFT2* | ADM92610.1 |
| *QsSVP1* | QSP002960.0 | *AtFT* | NP_176726.1 | *BvFLC* | ABN04205.1 |
| *QsSVP2* | QSP092511.0 | *AtSPL3* | AAM67271.1 | ***C. flexuosa*** | **NCBI accession** |
| *QsSVP3* | QSP072451.0 | *AtSPL5* | CAB56772.1 | *CfSPL9* | AGN29204.1 |
| *QsSVP4* | QSP018225.0 | *AtTFL1* | AAM27947.1 | *CfSOC1* | AGN29205.1 |
| ***P. trichocarpa*** | **NCBI accession** | *AtLEAFY* | AAM27931.1 | *CfFLC* | AGN29203.1 |
| *PtSPL* | XP_002309744.2 | ***M. domestica*** | **NCBI accession** | ***O. sativa* gene** | **NCBI accession** |
| *PtFT1* | EEE88631.1 | *MdSPL1* | ADL36827.1 | *OsHD3a* | BAO03040.1 |
| *PtFT2* | EEE88632.1 | *MdFT1* | BAI77730.1 | *OsSVP* | AAQ23144.2 |
| *PtSOC1* | AAP46287.1 | *MdSOC1* | NP_001280855.1 | *OsLEAFY* | AFA43522.1 |
| *PtSVP* | EEE90760.1 | *MdSVP* | AJW82922.1 | ***A. majus* gene** | **NCBI accession** |
| *PtFLC* | XP_011036097.1 | *MdLEAFY* | ABF84009.1 | *AmSBP1* | Q38741.1 |
| *PtLEAFY* | AAB51533.1 | ***P. persica*** | **NCBI accession** | ***A. chinensis*** | **NCBI accession** |
| ***H. vulgare*** | **NCBI accession** | *PpDAM1* | ABJ96361.2 | *AcSVP1* | AFA37967.1 |
| *HvSOC1* | AEX65782.1 | *PpDAM2* | ABJ96370.1 | *AcSVP2* | AFA37968.1 |
| *HvFLC* | ADI96238.1 | *PpDAM3* | ABJ96371.1 | *AcSVP3* | AFA37969.1 |
| ***E. grandis*** | **NCBI accession** | *PpDAM4* | ABJ96365.1 | *AcSVP4* | AFA37970.1 |
| *EgSOC1* | XP_010053874.1 | *PpDAM5* | ABJ96359.1 | ***M. truncatula*** | **NCBI accession** |
| ***A. alpina*** | **NCBI accession** | *PpDAM6* | ABJ96360.1 | *MtSVP1* | Medtr5g032520 |
| *AaSOC1* | AEH43352.1 | *PpLEAFY* | ABY78032.1 | *MtSVP2* | Medtr5g032150 |
| *AaPEP1* | FJ755930) | | | *MtSVP3* | Medtr5g066180 |

#### Supplementary Table II – List of primers

| Amplicon | Direction | sequence 5'-3' | Amplicon | Direction | sequence 5'-3' |
|---|---|---|---|---|---|
| ***QsFLC*** | Forward | GGAGTCCATAATGA GCCTTCA | ***QsSPL13*** | Forward | CCATTGCTTCGGAAAGTGAT |
| | Reverse | GGATGGGCCAACTG ATGAT | | Reverse | TGCACTAGGGAGGCTGAGT T |
| ***QsSOC1*** | Forward | GCAGCTGAAAATGC AAGGCT | ***QsDOF1*** | Forward | ACTTGTCGCCGTTATTGGAC |
| | Reverse | TTGCGCTTTGTTCTC CCTTCTG | | Reverse | GGTCATGATCGAAGCCAGTT |
| ***QsSVP1*** | Forward | GGACTTACCCGTGTG CTTGA | ***QsKINASE1*** | Forward | CCGGTTTGATTGGTTCACTT |
| | Reverse | ATGTCCGAGTCCAC AAGACC | | Reverse | ATTGCAAGCCCATTATCGAG |
| ***QsSVP2*** | Forward | TCGAATTCGCCAGCT CAAGT | ***QsCYP*** | Forward | CCCATCGTGTCCAAACTCTT |
| | Reverse | TGCTCAACATGGCGT AGGAG | | Reverse | AAtCTTCCTcCCTTCCTCCA |
| ***QsSVP3*** | Forward | ATGACGAGGAGGAA AATTCAGAT | ***QsS-LOCUS*** | Forward | AATTCGgTTGTTTGGTCAGC |
| | Reverse | TCATTTAGGAAAAG GTAGCCCC | | Reverse | tAaaGCGCCTGTTGAGACCT |
| ***QsSVP4*** | Forward | AATCTGGATTGAGC CGTGTG | ***QsWOX9*** | Forward | AGCCCATGGAGTaGgTGTTG |
| | Reverse | TCAGTGCCAATGTGT CTTCG | | Reverse | CCGaAAATCAAGAAGCAAG C |
| ***QsFT*** | Forward | CCTCTAGTTGTTGGG CGTGT | ***AtFT*** | Forward | CTAAGCTCTCAAGATCAAA GGCTTA |
| | Reverse | CCACCAATATCAAC CCTTGG | | Reverse | ACTAAAACGCAAAACGAAA GCGGTT |
| ***QsSPL4*** | Forward | ACCATAGGAGGCAC AAGGTG | ***QsACT*** | Forward | GCTGGATTCTGGTGATGGTG TGAGC |
| | Reverse | CGGCAACTCCTCTTT GTTTC | | Reverse | GCTTCAATGAGAGATGGCT GGAAGAGG |
| ***QsSPL9*** | Forward | ATGGAAATGGGTTC TGGCTCTC | ***QsPP2AA3*** | Forward | GGGTTCCCAACATCAAGTTC |
| | Reverse | TTAAAGTGACCAGT GCATCTGC | | Reverse | TGACCTGATCACTTGACTGC |
| ***QsLEAFY*** | Forward | ACTACCAACACCCTC GATGC | ***mIR156*** | Forward | CGTGACAGAAGAGAGTGAG AC |
| | Reverse | ACGTCCACCACTTTC CTTTG | ***mIR172*** | Forward | CGAGAATCTTGATGATGCTG CAT |
| **QsFLC gtw** | Forward | AAAAAGCAGGCTTA ACAATGGGGCGGAA GAAGGTGG | **QsFT gtw** | Forward | AAAAAGCAGGCTTAACAAT GCCCAGGGATAGGGATC |
| | Reverse | AGAAAGCTGGGTTC TATGGAAGCAAACT AAGTGTTTC | | Reverse | AGAAAGCTGGGTTTCATGTG GCAAAGCATTCAAG |
| **QsSVP1 gtw** | Forward | AAAAAGCAGGCTTA ACAATGGCGAGGGA GAAGATCAA | **QsSPL4 gtw** | Forward | AAAAAGCAGGCTTAACAAT GAAGCAACAGAAGGCGGTG |
| | Reverse | AGAAAGCTGGGTTT TAGAGAAGGGAAAG CCCTAA | | Reverse | AGAAAGCTGGGTTTTATCTG ATCTGGAAATGCTTGT |
| **QsSVP3 gtw** | Forward | AAAAAGCAGGCTTA ACAATGACGAGGAG GAAAATTCAGAT | **QsSPL9 gtw** | Forward | AAAAAGCAGGCTTAACAAT GGAAATGGGTTCTGGCTCT |
| | Reverse | AGAAAGCTGGGTTT CATTTAGGAAAAGG TAGCCCCAA | | Reverse | AGAAAGCTGGGTTTTAAAG TGACCAGTGCATCTGC |
| **QsSVP4 gtw** | Forward | AAAAAGCAGGCTTA ACAATGGCGAGAGA GAAGATTCAGA | **QsSPL13 gtw** | Forward | AAAAAGCAGGCTTAACAAT GAAAGCACCTTCTTGGGA |
| | Reverse | AGAAAGCTGGGTTT CACCCAGAGTAGGG TAGC | | Reverse | AGAAAGCTGGGTTCCCCAT GAGAAACATTCCCTG |
| **QsSOC1 gtw** | Forward | AAAAAGCAGGCTTA ACAATGGTGAGAGG AAAGACTCAG | | | |
| | Reverse | AGAAAGCTGGGTTG CGCTTTGTTCTCCCT TCTG | | | |
