## Supplementary figures and images for "Unisexual flower initiation in the monoecious *Quercus suber* L., a molecular approach"

### Supplemental figure 1

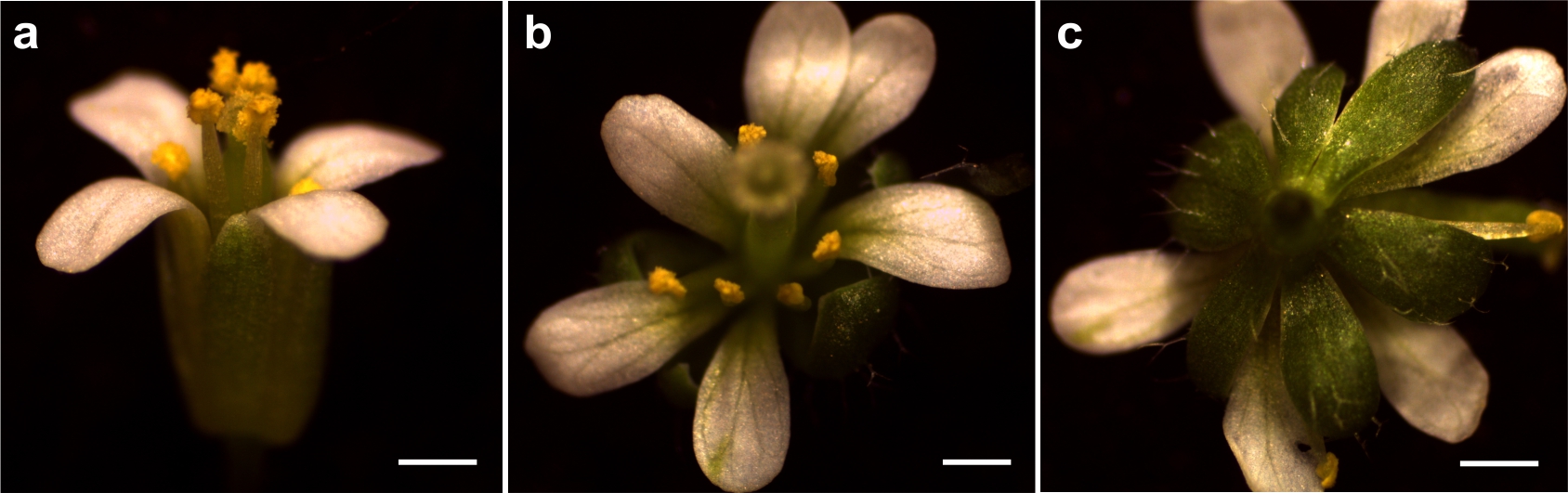

### Supplemental figure 2

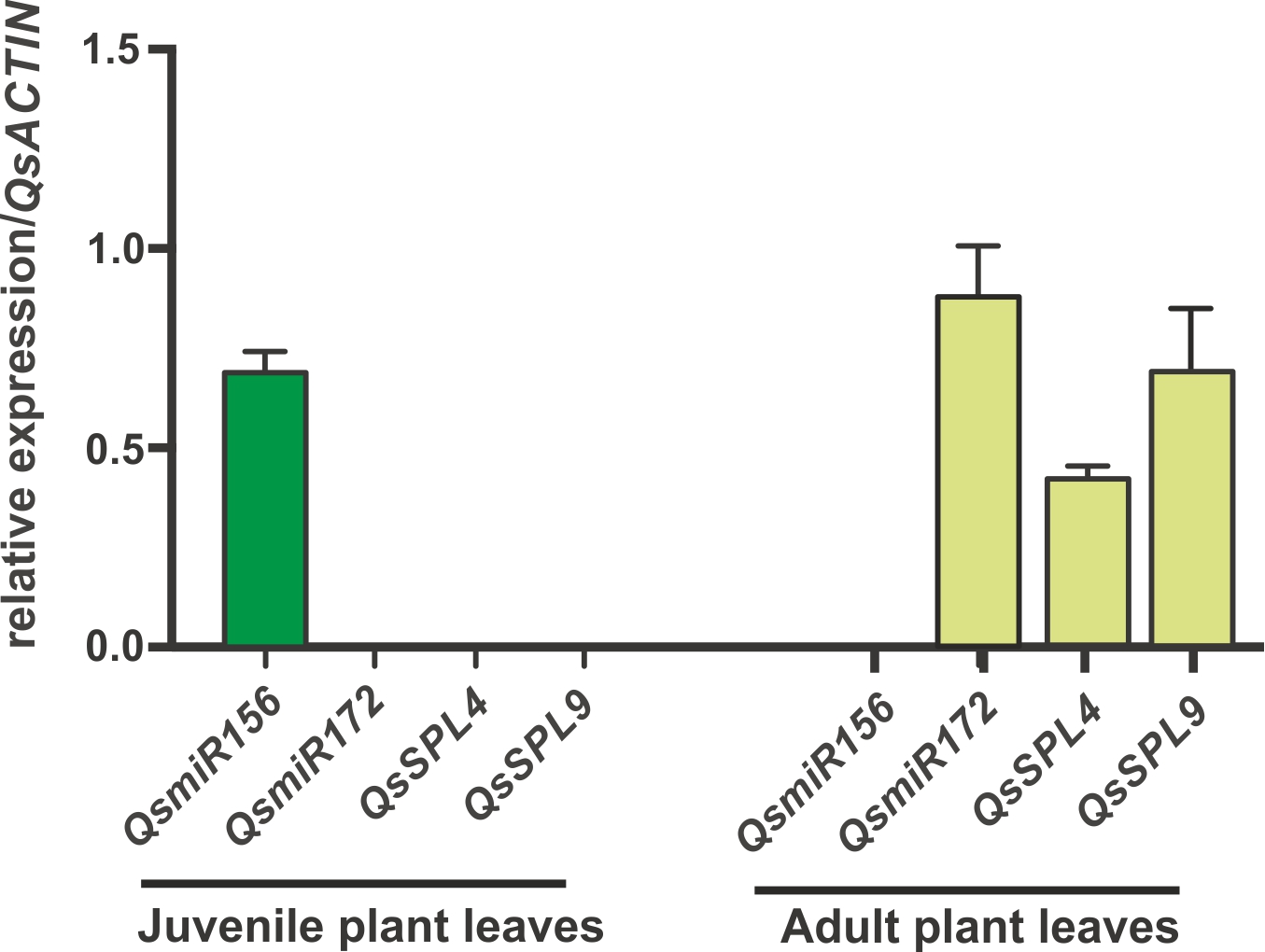

### Supplemental figure 3

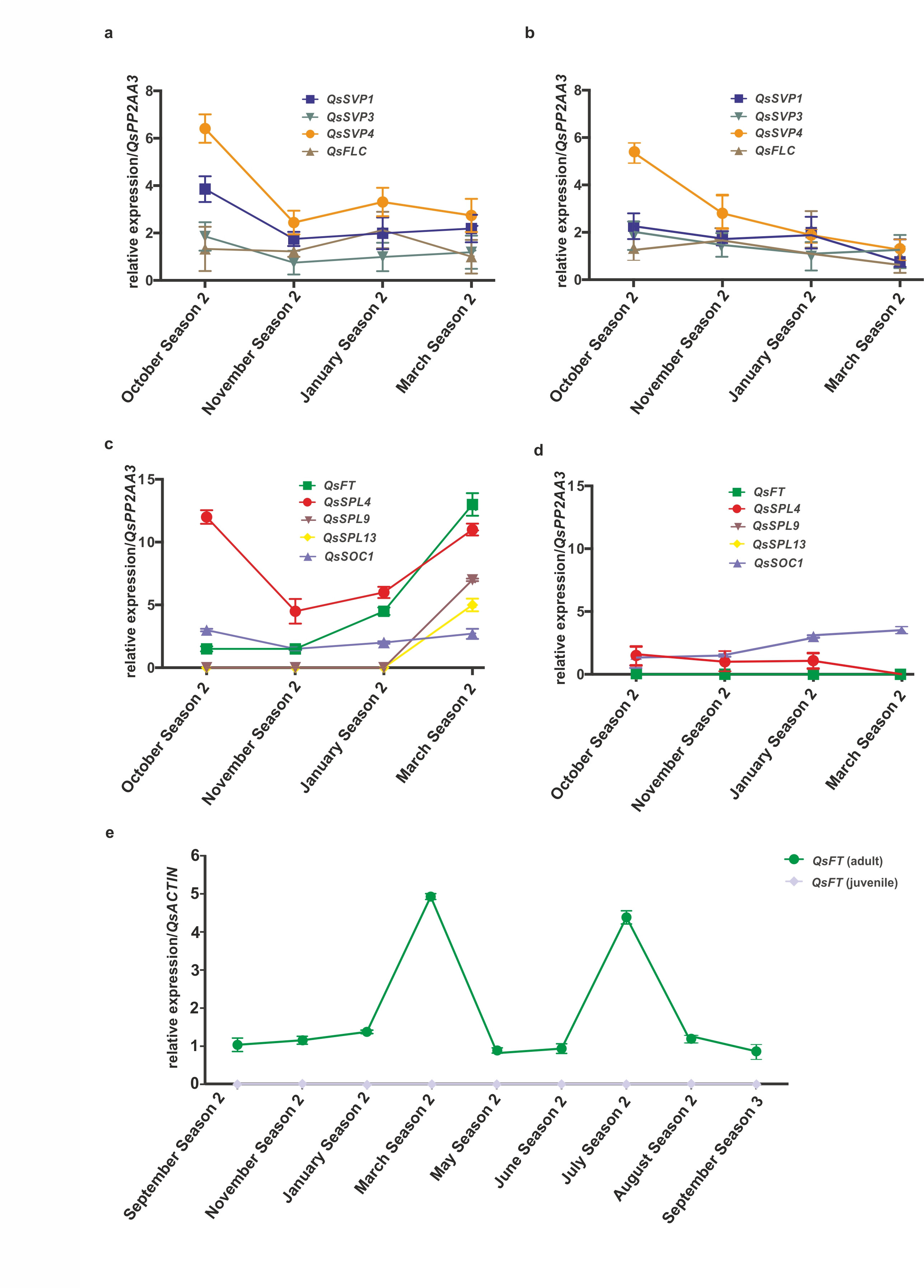

### Supplemental figure 4

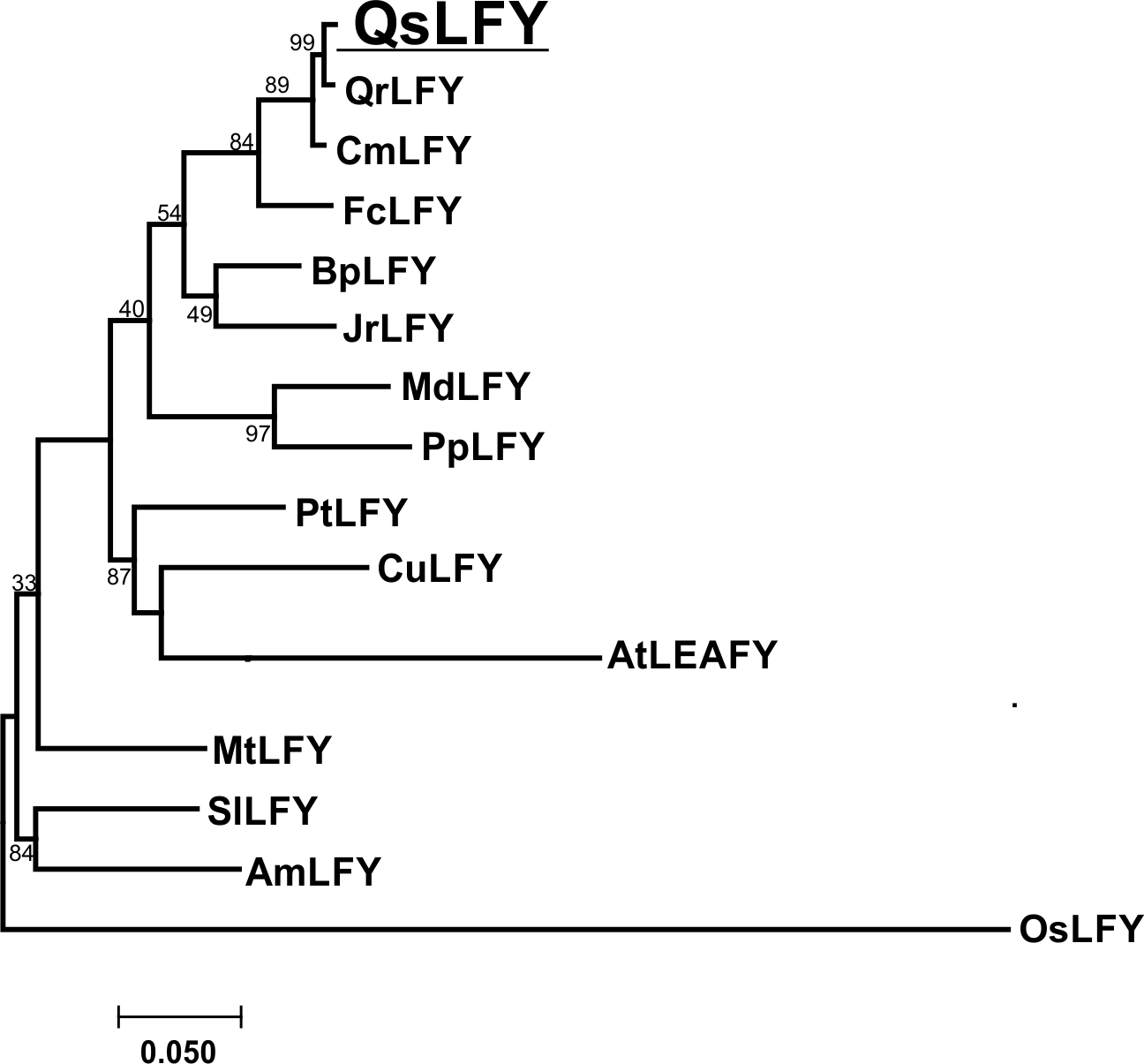

### Supplemental figure 5

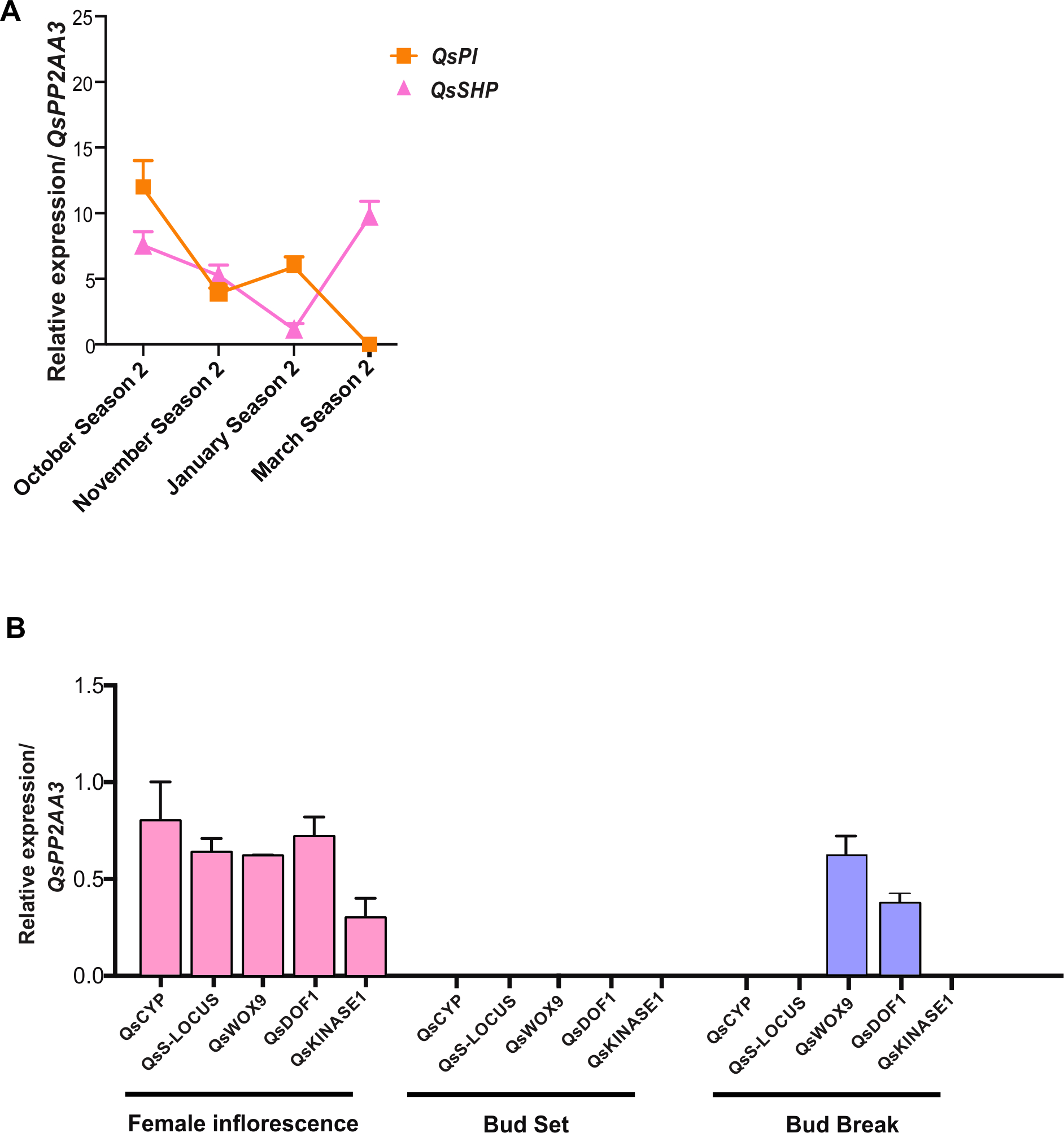
